# CRISPR PERSIST-On enables heritable and fine-tunable human gene activation

**DOI:** 10.1101/2024.04.26.590475

**Authors:** Y. Esther Tak, Jonathan Y. Hsu, Justine Shih, Hayley T. Schultz, Ivy T. Nguyen, Kin Chung Lam, Luca Pinello, J. Keith Joung

**Author notes:** These authors contributed equally (Y. Esther Tak, Jonathan Y. Hsu). These authors contributed equally (Justine Shih, Hayley T. Schultz).

## Abstract

Current technologies for upregulation of endogenous genes use targeted artificial transcriptional activators but stable gene activation requires persistent expression of these synthetic factors. Although general “hit-and-run” strategies exist for inducing long-term silencing of endogenous genes using targeted artificial transcriptional repressors, to our knowledge no equivalent approach for gene activation has been described to date. Here we show stable gene activation can be achieved by harnessing endogenous transcription factors (**EndoTF**s) that are normally expressed in human cells. Specifically, EndoTFs can be recruited to activate endogenous human genes of interest by using CRISPR-based gene editing to introduce EndoTF DNA binding motifs into a target gene promoter. This Precision Editing of Regulatory Sequences to Induce Stable Transcription-On (**PERSIST-On**) approach results in stable long-term gene activation, which we show is durable for at least five months. Using a high-throughput CRISPR prime editing pooled screening method, we also show that the magnitude of gene activation can be finely tuned either by using binding sites for different EndoTF or by introducing specific mutations within such sites. Our results delineate a generalizable framework for using PERSIST-On to induce heritable and fine-tunable gene activation in a hit-and-run fashion, thereby enabling a wide range of research and therapeutic applications that require long-term upregulation of a target gene.

## Results

Previously described “hit-and-run” methods for inducing stable repression of genes in mammalian cells rely on physiologic mechanisms used to maintain heritable silencing (e.g., DNA methylation)^1,2^. However, no DNA modifications are known that permit sustained gene activation and therefore we focused instead on leveraging the functions of activating EndoTFs that are already expressed in cells. To accomplish this, we envisioned that stable human gene activation might be achieved by using gene editing to introduce binding motifs for one or more activating EndoTFs into the promoter of a target gene (**Fig. 1a**, left side). These genetic edits would be permanent and would be expected to induce long-term sustained activation of that gene promoter, thereby bypassing the need for sustained expression of exogenous and/or synthetic transcription factors (**Fig 1a**, left panel). Because gene editors operate in a “hit-and-run” fashion, only transient expression of these components would be required to create the desired changes in promoter sequence and target gene expression. Thus, PERSIST-On differs from existing technologies for upregulation of genes such as CRISPR-based transcriptional activators (**CRISPRa**), which require continuous expression to mediate durable upregulation of the target gene (**Fig. 1a**, right panel).

**Figure 1.**
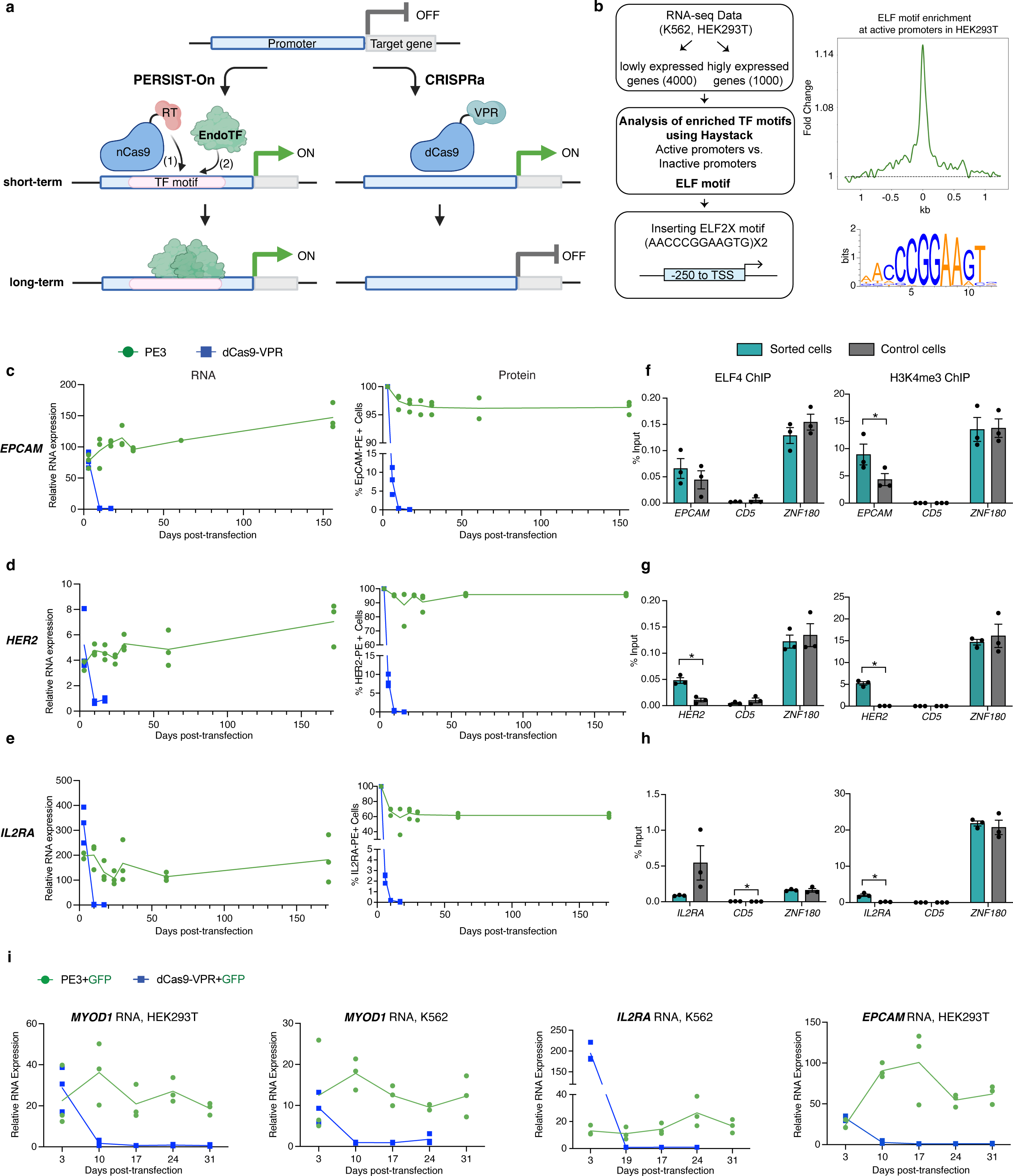
Stable and heritable gene activation by PERSIST-On with an ELFX2 motif compared with transient gene activation by CRISPRa. **a**, Schematics overview of long-term gene activation induced with PERSIST-On compared to short-term gene activation mediated by CRISPRa. With PERSIST-On, gene editing (shown here using prime editing) is used in the short-term (1) to insert a motif that is (2) recognized by an EndoTF expressed in the cell. Recruitment of EndoTF to the motif at the target promoter results in stable long-term target gene expression even after the gene editor is no longer expressed in the cell. By contrast, with CRISPRa, short-term expression of a synthetic activator (shown here using dCas9-VPR) leads to gene activation in the short-term. However, in the absence of long-term continued expression of the synthetic activator, target gene expression reverts to baseline levels. **b**, Computational workflow of TF motif screening that led to the selection of ELF motif sequence for insertion at the target promoters (left). Haystack analysis showing ELF motif enrichment at active promoters over inactive promoters in HEK293T cells (top right). Sequence logo of the ELF motif with height of each position indicating sequence conservation (bottom right). **c-e**, mRNA and cell-surface protein expression of the endogenous *EPCAM* gene in HEK293T cells or the endogenous *HER2* or *IL2RA* genes in K562 cells that had been sorted for cell-surface expression of the target gene product following transfection with plasmids expressing components for PERSIST-On (with the ELFX2 motif) or CRISPRa (with dCas9-VPR). Sorted edited cells were followed for expression over 156 days for *EPCAM* and 172 days for *HER2* and *IL2RA*. Unedited cells collected at various days were used as controls to establish baseline mRNA expression and to calculate fold-activation of RNA expression. Data shown are three biological replicates and the line represents the mean of those data over 156 days over 172 days. **f-h**, Chromatin immunoprecipitation (ChIP) experiments assessing binding of EndoTF ELF4 or presence of the H3K4me3 histone modification at the *EPCAM* promoter in sorted HEK293T cells (from **c**) or at the *IL2RA* or *HER2* promoters in sorted K562 cells (from **d-e)** that had been transfected with plasmids encoding components for PERSIST-On with the ELF2X motif. Control HEK293T or K562 cells were transfected with PE3 using non-targeting pegRNA and ngRNA. *CD5* is a negative control site and *ZNF180* is a positive control site for ELF4 binding in K562 and HEK293T cells. Closed circles indicate biological replicates (n=3), bars the mean of replicates, and error bars the s.e.m. * indicates significantly different from the control sample, p<0.05 (Student’s t-test, paired two-tailed test). **i**, Relative mRNA expression levels of the *MYOD1* gene in HEK293T and K562 cells, *IL2RA* gene in K562 cells, and *EPCAM* gene in HEK293T cells that were transfected with plasmids expressing components for PERSIST-On (with the ELFX2 motif) or CRISPRa (with dCas9-VPR) together with a GFP expression plasmid and then sorted for GFP expression. Sorted cells were followed for mRNA expression for 31 days. Data shown are three biological replicates and the line represents the mean of those data.

To identify a suitable EndoTF binding motif that might be used for an initial test of PERSIST-On in human cells, we searched for motifs that are more frequently found at the promoter regions of highly expressed genes compared to lowly expressed genes (**Fig. 1b**). We reasoned that these motifs might be bound by EndoTFs that are expressed and bound to active promoters in these cells. Using RNA-seq data, we performed a motif enrichment analysis^3^ contrasting the top 1000 highly expressed genes and bottom 4000 lowly expressed genes in HEK293T and K562 cells (**Fig. 1b**; **Online Methods**). This analysis yielded EndoTF motifs that are significantly enriched at highly expressed promoters in each cell line (**Supplementary Note 1**). Among the motifs commonly found in both cell lines, we chose the best 12 bp sequence based on motif models^4^ for the EndoTF ELF proteins (AACCCGGAAGTG), which were previously known to be recruited to constitutively active promoters^5^ (**Fig. 1b**). We created a dual direct repeat of this ELF motif (24 bps in length and hereafter referred to as the **ELF2X** motif) for use in subsequent experiments, reasoning that having two copies might increase the number of ELF EndoTFs bound when this sequence is inserted into an endogenous gene promoter (**Fig. 1b**).

We tested whether insertion of the ELF2X motif into the human *HER2*, *EPCAM*, and *IL2RA* promoters could lead to increases in expression of these genes in human cells. To accomplish this, we screened panels of CRISPR prime editing guide RNAs (**pegRNAs**) and nicking guide RNAs (**ngRNAs**) with prime editor 2 (PE2) protein^6^ in HEK293T cells for their abilities to insert the ELF2X motif at various distances up to 250 bps upstream of the transcription start site (**TSS**) (**Extended Data Figs. 1a – 1c**). Based on this initial screen, we chose four or five pegRNA/ngRNA pairs for each promoter (pink colored arrows, **Extended Data Figs. 1a – 1c**) and simultaneously assessed how efficiently they introduced the ELF2X motif (DNA editing activities) and how much they increased transcription of each target gene. We performed these experiments in cells in which each gene was not highly expressed: *EPCAM* in HEK293T cells and *HER2* and *IL2RA* in K562 cells. For each gene, we identified one or more pegRNA/gRNA pairs that could both mediate efficient insertion of the ELF2X motif and increase gene expression from the target promoter (**Extended Data Fig. 2**).

We next compared the efficiencies of PERSIST-On upregulation with ELF2X insertion to those induced by standard CRISPRa. To do this, we chose a single pegRNA/ngRNA combination for each gene that could be used to insert ELF2X into the *HER2* and *IL2RA* gene promoters in K562 cells and into the *EPCAM* promoter in HEK293T (black arrow, **Extended Data Fig. 2**). Because prime editing is relatively inefficient (**Extended Data Fig. 3a**), we transfected cells with plasmids encoding the prime editing components and then used flow sorting performed three days following transfection to enrich for cells with EpCAM, HER2, or IL2RA surface protein expression (**Extended Data Figs. 3a – 3c**, **Online Methods**). In parallel, we did the same with cells transfected with plasmids encoding components for CRISPRa (dCas9-VPR activator protein and a gRNA targeted to the same spacer sequence used by the pegRNAs to introduce the ELF2X motif) (**Online Methods**). As expected, with both ELF2X insertion and CRISPRa, all three endogenous genes showed increased RNA transcription (as judged by quantitative RT-PCR) relative to corresponding unedited (and unsorted) cell controls at day 3 post-transfection (**Figs. 1c – 1e; Extended Data Figs. 3b – 3c**). The initial magnitude of gene activation observed was comparable for the two approaches at the *HER2* and *EPCAM* genes and approximately two-fold higher with CRISPRa at the *IL2RA* gene (**Figs. 1c – 1e; Extended Data Figs. 3b – 3c**).

We additionally assessed the durability of activation induced by ELF2X insertion and CRISPRa by quantifying activation of *EPCAM*, *IL2RA*, and *HER2* over a longer timeframe. Because we used transient transfection to introduce the plasmids encoding components for prime editing or CRISPRa into cells, we expected that these episomal DNAs will be lost rapidly as the cells divide due to degradation and dilution. Notably, with PERSIST-On, the transcriptional activation of target gene mRNA observed for all three targets was stable for at least nearly six months (170 days) (**Figs. 1c – 1e**). Consistent with this effect being due to the ELF2X insertion, we observed that the percentages of DNA alleles harboring that insertion were also stable over that same timeframe in these edited cells (**Extended Data Fig. 4)**. By contrast, with CRISPRa, the increased target gene mRNA levels more rapidly declined back to baseline within ten days (**Figs. 1c – 1e**), presumably due to loss of the plasmids encoding (and therefore the expression of) the dCas9-VPR and gRNAs needed to maintain activation. These same differential patterns of stable and transient gene activation were also observed at the protein level using flow cytometry with cell surface staining to assess expression of the EpCAM, IL2RA, or HER2 proteins (**Figs. 1c – 1e**). We did observe an initial decrease in the percentage of IL2RA-positive cells with ELF2X insertion, which we speculate is most likely due to false positive staining of cells in the initial sorted sample (although we cannot rule out toxicity in a subpopulation due to prime editing). ChIP-qPCR analysis performed on the cells that underwent PERSIST-On showed increased H3K4me3 deposition at all three gene promoters consistent with increased transcription (**Figs. 1f – 1h**). To attempt to determine the identity of the EndoTF that binds to the ELF2X motif, we used ChIP-qPCR to test whether the endogenous ELF4 transcription factor (which is highly expressed in HEK293T and K562 cells) might be the EndoTF that binds to the ELF2X motif. We detected significant enriched binding of ELF4 at the *HER2* promoter but not at the *EPCAM* and *IL2RA* promoter (**Figs. 1f – 1h**), suggesting that a different member of the ELF transcription factor family might bind to the ELF2X insertion at these latter promoters.

Because the experiments described above used flow sorting to enrich for high expression of the target gene product, we sought to test a more generalizable strategy to use PERSIST-On that does not require the target gene to be cell surface expressed. For this alternative approach, we included a GFP expression plasmid in our transfections and then used flow sorting to select for cells that had been most efficiently transfected with plasmids encoding PERSIST-On or CRISPRa components rather than for cell surface expression of the target gene product (**Online Methods**). Using this strategy, we inserted ELF2X into the *EpCAM* promoter in HEK293T cells and the *IL2RA* promoter in K562 cells (**Extended Data Fig. 3d**). We also inserted ELF2X into the *MYOD1* (a non-surface-expressed gene) promoter in both HEK293T and K562 (**Extended Data Fig. 3d**). We additionally performed matched transfections with plasmids encoding components needed for CRISPRa-mediated activation of these same promoters (using gRNAs targeting the same spacer sequences as the pegRNAs used in the PERSIST-On experiments). Immediately following sorting on post-transfection day 3, we observed transcriptional activation (as measured by quantitative RT-PCR) with PERSIST-On and CRISPRa for all three genes (**Fig. 1i**). As expected, because we did not sort directly for target gene expression, we saw somewhat lower but still robust levels of activation for *EpCAM* and *IL2RA* than the earlier experiments shown in **Figs. 1c and 1e**, respectively (compare **Extended Data Figs. 3b and 3e**). Importantly, we again observed that gene activation in all four cases was stable out to 31 days with ELF2X insertion but returned to baseline by day 10 with CRISPRa (**Fig. 1i**). The percentage of genomic DNA alleles bearing the targeted ELF2X insertion also remained stable out to 31 days in all cases (**Extended Data Fig. 5)**. Taken together, these data across four different endogenous target genes and two cell types demonstrate that PERSIST-On can provide a general strategy for stable activation of gene expression.

Having established the efficacy of PERSIST-On with the ELF2X motif, we next sought to determine whether the method might also work with other EndoTFs and their associated binding motifs. To test this, we first compiled a list of consensus sequences from a set of 286 archetype EndoTF motifs, obtained by clustering and consolidation of similar TF motif models^7^ (**Online Methods**) and then multimerized each of these motifs to create 286 final sequences that were each 24 bps in length (**Supplementary Table 1**). We designed pegRNAs to insert each of these 286 sequences as well as five negative control sequences (also all 24 bps in length) into the same locations at which we had previously inserted the ELF2X motif in the *IL2RA* and *HER2* promoters (**Supplementary Table 2**). We then used the Multiplexing Of Site-specific

Alterations for *In situ* Characterization (**MOSAIC**) method (https://doi.org/10.1101/2024.04.25.591078) to rapidly screen these pegRNAs in pooled format to determine which of the 286 archetype EndoTF motifs might activate the *IL2RA* and *HER2* target promoters in K562 cells (**Fig. 2a**; **Online Methods**). Three days after transfection of EndoTF pegRNA libraries with prime editing components (**Online Methods**), we used flow sorting and targeted amplicon sequencing to determine which specific archetype motifs were enriched in the HER2- and IL2RA-positive populations relative to the HER2- and IL2RA-negative populations, respectively (**Fig. 2a**; **Online Methods**). For both gene promoters, this yielded multiple archetype motifs whose insertions were associated with enrichment (presumably due to target gene activation) at similar or higher levels compared with those observed with the previously validated ELF2X motif (**Figs. 2b - 2c**). Many of the motifs we identified were enriched with both the *IL2RA* and *HER2* promoters or not enriched with either promoter (**Fig. 2d**). Interestingly, many of these activating archetype motifs are for transcription factor families characterized as pioneer transcription factors capable of directly binding to and opening inaccessible target DNA and recruiting other transcription factors and cofactors for gene activation^8^. We individually tested nine motifs that had been enriched in the MOSAIC IL2RA experiments for their abilities to activate the *IL2RA* promoter in K562 cells and found that all up-regulated the gene (**Fig. 2e**). The mean activation levels observed with these nine motifs ranged from 2.8- to 161-fold even with only modest frequencies of motif insertion ranging 1 to 20% (**Fig. 2e**). The CREB/ATF/3-2X motif induced the highest fold-activation and therefore we used this sequence in additional experiments described below.

**Figure 2:**
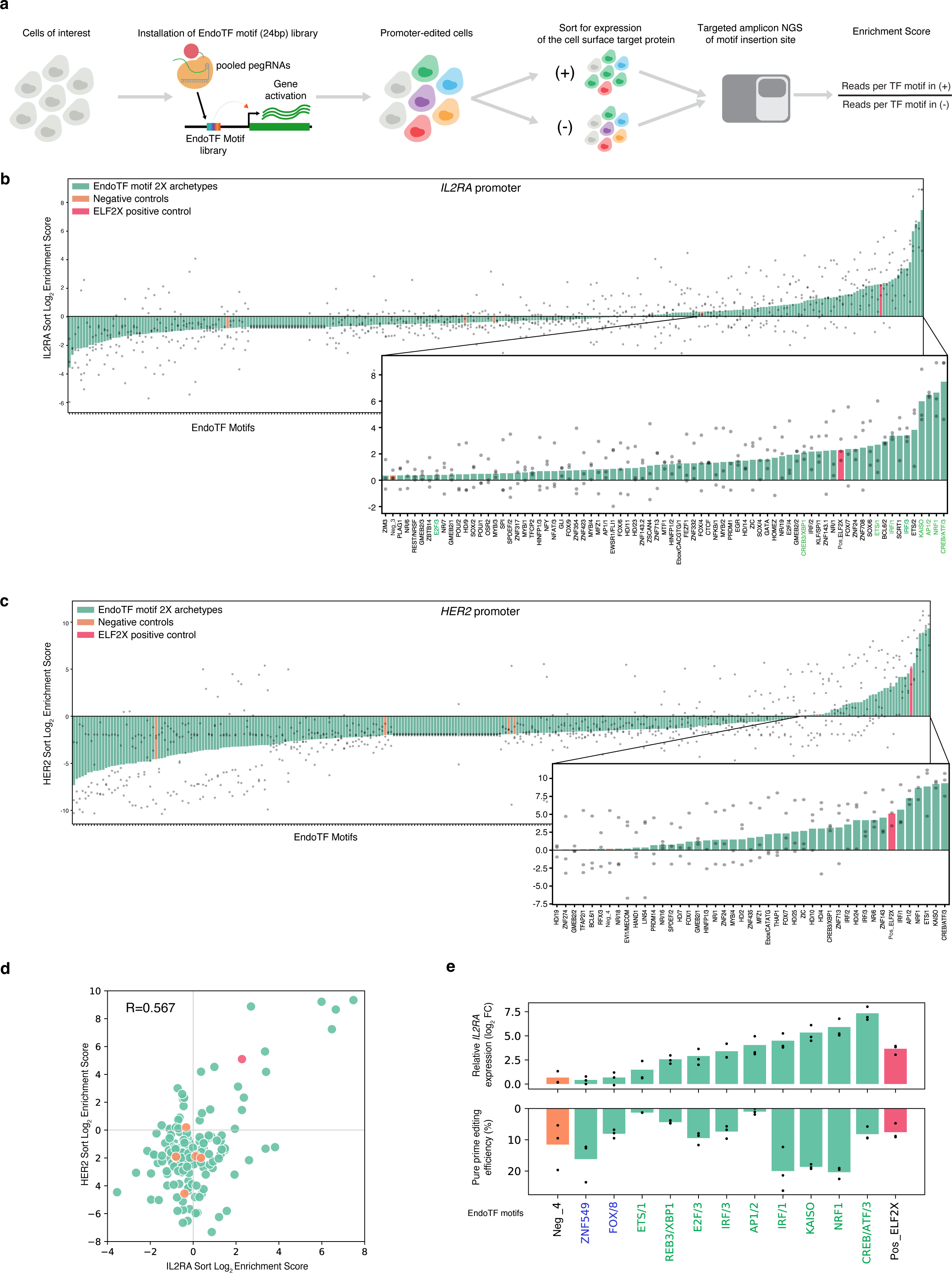
Large-scale MOSAIC-based screening of various EndoTF archetype motifs for mediating PERSIST-On in human cells. **a**, Schematic overview of a MOSAIC-based high-throughput pooled screening strategy to test various EndoTF motifs for their PERSIST-On activities. A pool of pegRNAs designed to install a library of EndoTF archetype motifs, the ELF2X motif (as a positive control), and five random DNA sequences (as negative controls) is used with MOSAIC to install these sequences into an endogenous gene promoter in human cells. Flow cytometry is then used to sort cells based on presence (+) or absence (-) of target gene product expression on the cell surface. Enrichment scores can then be calculated for each EndoTF motif by using the ratio of the number of alleles with the motif in the (+) cells over the number of motif alleles in the (-) cells (as determined by targeted amplicon sequencing). **b**, Enrichment scores of 286 EndoTF motifs (green bars), the positive control ELF2X motif (red bar), and five negative control sequences (orange bars) introduced into the *IL2RA* promoter in K562 cells using the MOSAIC-based strategy described in **a**. Sequences are arranged along the x-axis by their mean enrichment scores (bars) based on three biological replicates (grey circles). An expanded view of the graph shows the identities of sequences with positive mean enrichment scores greater than the highest scoring negative control. High-scoring archetypal TF motifs selected for individual validation (see **e** below) are highlighted in green text. Neg: negative control, Pos: positive control. **c**, Enrichment scores of 286 EndoTF motifs (green bars), the positive control ELF2X motif (red bar), and five negative control sequences (orange bars) introduced into the *HER2* promoter in K562 cells using the MOSAIC-based strategy described in **a**. Data are depicted in the same fashion as those in **b**. **d**, Scatterplot showing the mean enrichment scores of EndoTF motifs (from the experiments of **b** and **c** above) for the *IL2RA* (x-axis) and *HER2* (y-axis) promoters. Green dots are from 189 archetype EndoTF motifs that were selected based on the number of reads from NGS (>100 total reads per motif), orange dots are the negative controls, and the red dot is the positive control (ELF2X motif). Correlation coefficient (R) shown was determined with Spearman’s Rho. **e**, Individual validation of nine high-scoring archetypal TF motifs (names shown in green text) and two low-scoring archetypal TF motifs (names shown in blue text) from **b** inserted at the *IL2RA* promoter in K562 cells using CRISPR prime editing. Top panel shows fold-activation of *IL2RA* gene expression observed and bottom panel shows the efficiency of motif introduction in the same populations of transfected cells. Circles indicate three biological replicates and bars represent the mean of those data. Neg: negative control, Pos: positive control.

We next explored whether systematically mutating archetype motifs capable of causing activation might provide a means to fine tune the magnitude of gene upregulation induced by their insertion into a target promoter. We tested this strategy by constructing MOSAIC pegRNA libraries designed to introduce a series of mutated ELF2X or CREB/ATF/3-2X sequences harboring each of the three possible base substitutions at each position within both copies of each of these activating archetype motifs (**Supplementary Table 1**). We then used MOSAIC to introduce these sequences (and five negative control sequences) into the *HER2* and *IL2RA* promoters in K562 cells (**Fig. 3a**). Three days after transfection, we processed these cells in two different ways: (1) we used flow to cytometry to sort some of the cells into populations that were positive or negative for HER2 or IL2RA cell surface protein expression and (2) we performed chromatin immunoprecipitation (ChIP) on the remaining unsorted cells using an antibody against either the ELF4 or ATF4 transcription factors we expected would bind to the ELF2X or CREB/ATF/3-2X sequences, respectively (the processing of these ChIP experiments is described in the next paragraph). We performed targeted amplicon sequencing of the insertion site on the sorted and unsorted cells to identify which mutated motif sequences were enriched in cells with high target gene expression (**Fig. 3a**). For both the mutated ELF2X and CREB/ATF/3-2X sequences, we observed that a small subset of these sites (and the original unmutated sites) was enriched in cells with expression of IL2RA or HER2 (ranges of 2- to 4-fold or 2- to 256-fold enrichment, respectively) whereas the negative controls and other mutated sequences were not enriched (**Figs. 3b – 3c; Extended Data Figs. 6a – 6b**). Notably, we observed a high degree of correlation in enrichment of each set of mutated sequences with the two different target genes (**Figs. 3d – 3e**) suggesting consistency in the effects of these various mutated motifs on target gene activation. We also confirmed that a range of mutated ELF2X and CREB/ATF/3-2X sequences enriched in our MOSAIC experiments could each induce various levels of IL2RA activation when tested individually (**Extended Data Figs. 6c – 6d**).

**Figure 3:**
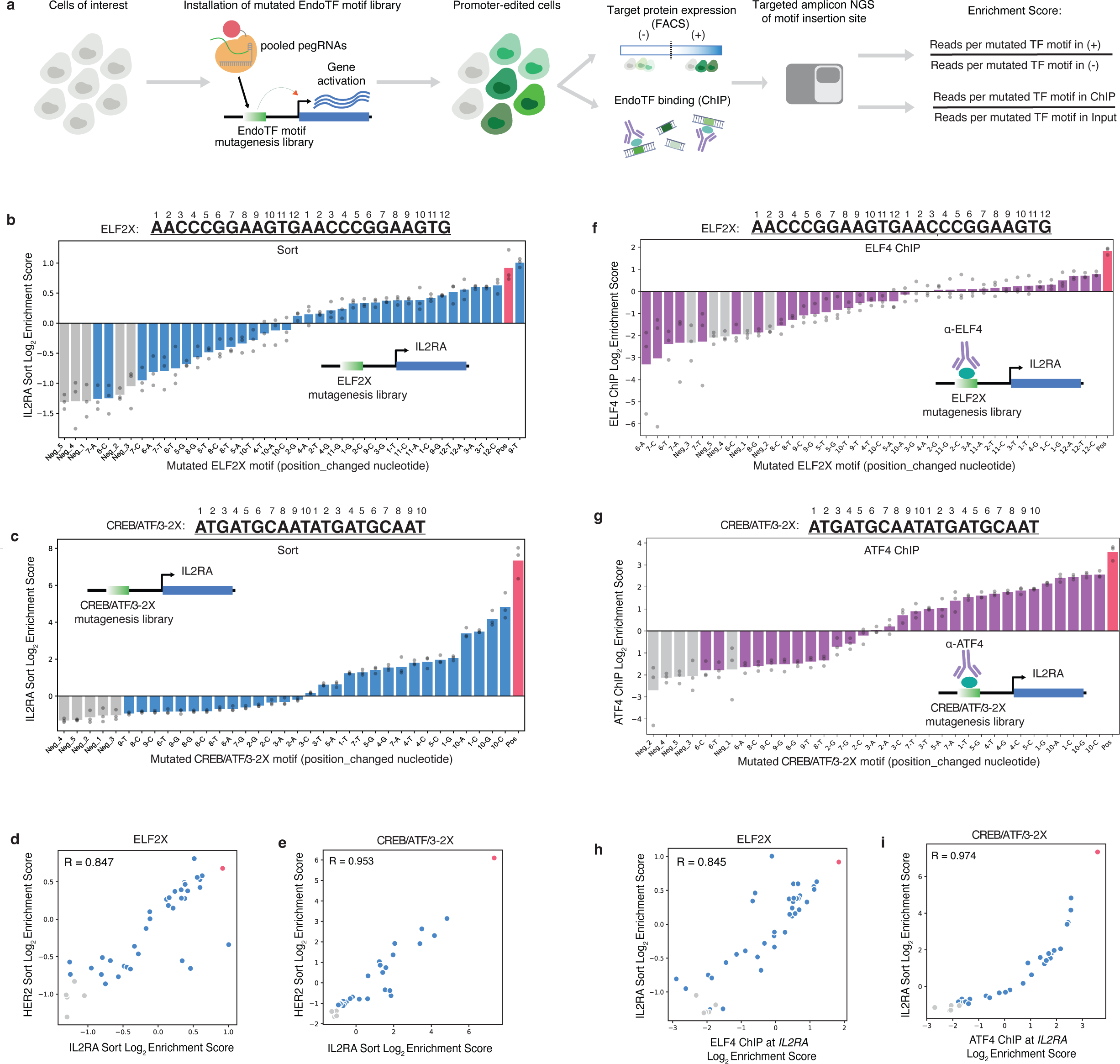
Fine-tuning of PERSIST-On activation of the *IL2RA* gene promoter by MOSAIC-based large-scale screening of systematically mutated EndoTF motifs. **a**, Schematic overview of a MOSAIC-based high-throughput pooled screening strategy to test systematically mutated EndoTF motifs for their PERSIST-On activities. A pool of pegRNAs designed to install a library of mutated EndoTF archetype motifs, the original unmutated motif, (as a positive control), and five random DNA sequences (as negative controls) is used with MOSAIC to install these sequences into an endogenous gene promoter in human cells. A portion of edited cells is then sorted into populations based on presence (+) or absence (-) of target gene product expression on the cell surface. Enrichment scores for these sorted cells can then be calculated for each motif introduced by using the ratio of the number of alleles with the motif in the (+) cells over the number of motif alleles in the (-) cells (as determined by targeted amplicon sequencing). The other portion of cells is used to perform ChIP using an antibody against an EndoTF that binds to the original unmutated motif. Enrichment scores for these ChIP experiments can then be calculated for each motif introduced by using the ratio of the number of alleles with the motif in the immunoprecipitated genomic DNA over the number of motif alleles in the original input genomic DNA (prior to ChIP) (as determined by targeted amplicon sequencing). **b-c**, Enrichment scores of a large series of systematically mutated ELF2X (**b**) or CREB/ATF/3-2X (**c**) motifs (blue bars) inserted into the *IL2RA* promoter in K562 cells that had been flow sorted based on IL2RA surface expression as illustrated in (**a**). Mutated motifs are denoted by “base position -base changed nucleotide”. Enrichment scores are also shown for the original unmutated ELF2X or CREB/ATF/3-2X motifs (red bars) and five random negative control DNA sequences (grey bars). Closed circles indicate biological replicates (n=3) and bars represent the means of these replicates. **d-e**, Scatterplot showing mean enrichment scores (from cells flow sorted for cell surface IL2RA expression) of various mutated ELF2X (**d**) or CREB/ATF/3-2X (**e**) motifs at the *IL2RA* (x-axis) and *HER2* (y-axis) promoters (data from **b**, **c**, and **Extended Data** Figs. 6a **-6b**). Correlation coefficients (R) were determined with Spearman’s Rho. Blue dots: mutated ELF2X or CREB/ATF/3-2X motifs; grey dots: negative controls, red dots: unmutated ELF2X or CREB/ATF/3-2X motif. **f-g**, Enrichment scores of a large series of systematically mutated ELF2X (**f**) or CREB/ATF/3-2X (**g**) motifs (purple bars) inserted into the *IL2RA* promoter in K562 cells that had been analyzed by ChIP as illustrated in (**a**). Mutated motifs are denoted by “base position - base changed nucleotide”. Enrichment scores are also shown for the original unmutated ELF2X or CREB/ATF/3-2X motifs (red bars) and five random negative control DNA sequences (grey bars). Closed circles indicate biological replicates (n=3) and bars represent the means of these replicates. **h-i**, Scatterplots showing ChIP-based enrichment scores (x-axes) and IL2RA surface expression-based (flow sorting) enrichment scores (y-axes) for motifs associated with ELF4 (**h**) or ATF4 (**i**) EndoTFs. Correlation coefficients (R) were determined with Spearman’s Rho. Blue dots: mutated ELF2X or CREB/ATF/3-2X motifs; grey dots: negative controls, red dots: unmutated ELF2X or CREB/ATF/3-2X motif.

Having confirmed that mutating archetype motifs could produce variable levels of gene activation, we next examined whether these effects might be associated with altered binding efficiencies of EndoTFs to those mutated sequences. To do this, we used targeted amplicon sequencing to ascertain which mutated motifs were enriched in the K562 ChIP experiments described in (2) in the paragraph above. These analyses revealed that a small subset of mutated ELF2X and CREB/ATF/3-2X sites (and the original unmutated ELF2X and CREB/ATF/3-2X sites) were positively enriched while the negative controls and the remaining other mutated sequences were not (**Figs. 3f – 3g; Extended Data Figs. 6e – 6f**). Notably, scatterplots showed a high degree of correlation (Spearman’s correlation coefficient (R) values ranging from 0.85 to 0.96) between enrichment scores of sites in the ChIP experiments and enrichment scores of those sites in the same K562 cells that we had sorted for target gene expression (**Figs. 3d – 3e; Extended Data Figs. 6g – 6h**), consistent with the idea that the degree of archetype motif occupancy likely impacts the magnitude of gene activation. Interestingly, the shape of data points in the scatterplots show a more sigmoidal shape for the CREB/ATF/3-2X sites than for the ELF2x sites with both target genes, perhaps reflecting cooperative DNA binding of the heterodimeric CREB/ATF EndoTF complex compared with the presumably non-cooperative binding of monomeric ELF EndoTF to these sites (**Figs. 3h – 3i; Extended Data Figs. 6i – 6j**). Our results collectively suggest that the degree of DNA binding of EndoTFs and thereby the magnitude of target gene promoter activity can be modulated by introducing mutations into the archetype motifs used in PERSIST-On.

## DISCUSSION

PERSIST-On provides a generalizable strategy to induce stable activation of endogenous genes that can be flexibly tuned in different ways to achieve desired levels of expression. Although our studies were performed in human cells, because EndoTFs are used by and expressed in all organisms, PERSIST-On should in principle be generalizable to any cell into which gene editing components can be efficiently delivered and for which the binding sites of EndoTFs are known. Our experiments show that the method requires assessment and optimization of various parameters: (1) choice of EndoTF binding motif(s) used, which can be selected based on EndoTFs expressed in the cell type of interest and/or empirical testing of various different motifs using MOSAIC to identify the optimal choice; (2) location of the insertion site of the EndoTF motif within the target gene promoter, both because the distance from the TSS may influence the degree of activation and because the efficiency of editing to achieve insertion may differ from site to site; and (3) introduction of mutations into the archetype motif to vary the level of activation induced, a parameter that can also be assessed using MOSAIC. These different variables can be adjusted and exploited to achieve the desired level of target gene expression.

Other parameters might also be varied to modulate the level of target gene expression that we did not test in our studies here. These include the degree of archetype multimerization (e.g., we used only two copies for most of our experiments but could use more), the length of the spacing between individual motifs within a multimerized sequence, and the use of heteromeric combinations of motifs, all of which can be deployed to leverage transcriptional activator synergy. We further note that it might be of interest to determine if the simultaneous insertion of archetype motifs into not just a promoter but also into an associated long-range element (e.g., one that functions as a transcriptional enhancer in other cell types) might lead to synergistic activation similar to what we recently observed when simultaneously directing artificial CRISPR-based activators to those two types of sequences^9^. An additional parameter that could impact the magnitude of gene activation and that might also be leveraged for therapeutic and other applications is cell-type-specific effects. For example, one can imagine that the presence and binding of other cell-type-specific EndoTFs to the target promoter might influence the degree of gene activation that occurs when an archetype motif is inserted ectopically into that same sequence. In addition, one can also envision that insertion of archetype motifs for cell-type-specific TFs might permit gene activation to only occur in the cells in which those EndoTFs are expressed, providing an additional means to achieve cell-type specificity beyond restricting or limiting what tissues or cells PERSIST-On components are delivered into.

We also note that the MOSAIC-based strategy we used to systematically mutate base positions within ectopically inserted EndoTF binding sites can provide a much richer profile of EndoTF DNA-binding specificity than is possible with standard genome-wide ChIP assays. This is because our MOSAIC-based approach comprehensively assesses all possible base substitutions at every position within an EndoTF binding site, thereby enabling the generation of position weight matrices (**PWM**s) and sequence logos with not only positive base preferences at each position but also negative preferences as well. For example, we used flow sorted cells for IL2RA and HER2 expression and ChIP data for binding of ELF4 and ATF4 to their ectopic inserted sites at the *IL2RA* and *HER2* promoters to generate PWMs and sequence logos showing positive and negative base preferences at every position within the ELF2X and CREB/ATF/3-2X sites (**Extended Data** Fig. 7). A similar MOSAIC-based strategy could also be deployed to define the binding specificities of EndoTFs to their normal binding sites in any given promoter. Doing so could provide a more physiologic study of EndoTF binding activities *in situ* in their native chromatin contexts as opposed to previously described *in vitro* analysis of binding specificities of EndoTFs in the literature that are performed in synthetic contexts^10,11^.

Finally, we envision that PERSIST-On may provide a potentially promising strategy to explore in future studies for therapeutic applications requiring stable long-term activation of target gene expression. Because PERSIST-On uses gene editing, a widely used technology already being advanced into the clinic, it can leverage all existing and future viral and non-viral platforms used for delivery of these components into various human cells and tissues. Although we used prime editing to introduce EndoTF motifs in this study, one can readily envision that other gene editing methods such as targeted nucleases (to induce synthetic double-stranded DNA capture^12^ or homology directed repair) or base editors could also be used for PERSIST-On. Lastly, because PERSIST-On introduces edits to *cis*-acting sequences, this strategy should achieve a more consistent level of gene activation relative to other strategies that introduce synthetic *trans*-acting synthetic transcriptional activators (e.g., CRISPRa), where it can be more challenging to not only maintain expression of these factors but also to control their levels of expression from cell to cell. This could be of particular importance when treating disorders of haploinsufficiency where the desired therapeutic window for restoring wild-type gene activation may be particularly narrow^13^. Consistency of gene activation may also be critical for many other therapeutic and research applications using gene activation given recent work predicting that over 8% of human coding genes are likely to be triplosensitive^13^. Taken together, our work provides an important initial proof-of-concept for the PERSIST-On strategy that should motivate further studies to explore the therapeutic utility of this technology.

## EXTENDED DATA FIGURE LEGENDS

**Extended Data Figure 1. Screening for pegRNAs that efficiently insert the ELF2X motif at the *HER2*, *IL2RA*, or *EPCAM* promoter in HEK293T cells.**

**a-c**, Genomic locations of 24 pegRNAs designed to insert the ELF2X motif at various locations within the *HER2* (**a**), *IL2RA* (**b**), or *EPCAM* (**c**) promoter in HEK293T cells (left). Arrows indicate direction of gene transcription. Frequencies of ELF2X motif insertion (y-axes) using 24 pegRNAs with the PE3 strategy at the *HER2*, *IL2RA* or *EPCAM* promoter in HEK293T cells (right). Pink arrows indicate the five best performing pegRNA + ngRNA combinations selected for subsequent testing for effects on target gene expression (shown in **Extended Data** Fig. 2). Open circles indicate biological replicates (n = 2 for *IL2RA* or n = 3 for *HER2* and *EPCAM*), bars represent the mean of those replicates, and error bars represent the s.e.m.

**Extended Data Figure 2. Activation of gene expression induced by ELF2X motif insertion at the *EPCAM, IL2RA*, or *HER2* promoter with various high efficiency pegRNAs.**

Five pegRNAs with the highest prime editing activities for inserting the ELF2X motif at the *EPCAM*, *IL2RA* or *HER2* promoter (identified in the experiments shown in **Extended Data Fig. 1**) were renamed (1-5) and re-tested for their ELF2X insertion efficiencies at the *EPCAM* promoter in HEK293T cells and the *IL2RA* or *HER2* promoter in K562 cells (left graphs). These same cells were also tested for fold-activation of the target gene mRNA (right graphs). Closed circles indicate biological replicates (n = 3), bars represent the means of those replicates, and error bars are the s.e.m. Arrows indicate the pegRNA + ngRNA combinations subsequently used for each of these genes in the experiments of **Figs. 1c - 1e**.

**Extended Data Figure 3. Prime editing and target gene activation efficiencies in unsorted and sorted PERSIST-On (with ELF2X insertion) and CRISPRa (with dCas9-VPR) cells.**

**a**, Efficiencies of ELF2X insertion in cells that were modified by PERSIST-On (from **Figs. 1c – 1e**) before and after sorting (three days post-transfection) for surface expression of the indicated target gene product. ELF2X was inserted into the *IL2RA* and *HER2* promoters in K562 cells and into the *EPCAM* promoter in HEK293T cells. Open circles indicate biological replicates (n= 3), bars represent the mean of those replicates, and error bars are the s.e.m. for those replicates.

**b**, Fold-activation of target gene RNA expression in the same cells shown in **a**. Open circles indicate biological replicates (n= 3), bars represent the mean of those replicates, and error bars are the s.e.m. for those replicates.

**c**, Fold-activation of target gene RNA expression in cells that were modified by CRISPRa (from **Figs. 1c – 1e**) before and after sorting (three days post-transfection) for surface expression of the indicated target gene product. dCas9-VPR together with a gRNA targeted to the same spacer sequence targeted by the ELF2X insertion were co-expressed in K562 cells (for the *IL2RA* and *HER2* promoters) and in HEK293T cells (for the *EPCAM* promoter). Open circles indicate biological replicates (n= 3), bars represent the mean of those replicates, and error bars are the s.e.m. for those replicates.

**d**, Efficiencies of ELF2X insertion in cells that were modified by PERSIST-On (from **Figs. 1c – 1e**) before and after sorting (three days post-transfection) for expression of GFP from a co-transfected plasmid. ELF2X was inserted into the *IL2RA* and *HER2* promoters in K562 cells and into the *EPCAM* promoter in HEK293T cells. Open circles indicate biological replicates (n= 3), bars represent the mean of those replicates, and error bars are the s.e.m. for those replicates. **e**, Fold-activation of target gene RNA expression in the same cells shown in **d**. Open circles indicate biological replicates (n= 3), bars represent the mean of those replicates, and error bars are the s.e.m. for those replicates.

**f**, Fold-activation of target gene RNA expression in cells that were modified by CRISPRa (from **Figs. 1c – 1e**) before and after sorting (three days post-transfection) for expression of GFP from a co-transfected plasmid. dCas9-VPR together with a gRNA targeted to the same spacer sequence targeted by the ELF2X insertion were co-expressed in K562 cells (for the *IL2RA* and *HER2* promoters) and in HEK293T cells (for the *EPCAM* promoter). Open circles indicate biological replicates (n= 3), bars represent the mean of those replicates, and error bars are the s.e.m. for those replicates.

**Extended Data Figure 4. Stability of ELF2X motif insertion at target promoters in cells modified by PERSIST-On and sorted for surface expression of the target gene**

Efficiencies of ELF2X insertion in cells that were modified by PERSIST-On (from **Figs. 1c – 1e**) and sorted for surface expression of the indicated target gene product were assessed at various time points post-transfection. Insertion frequencies at the *IL2RA* and *HER2* promoters were assessed in K562 cells for 170 days post-transfection and at *EPCAM* promoter in HEK293T cells for 150 days post-transfection. Data shown are three biological replicates and the line represents the mean of those data.

**Extended Data Figure 5. Stability of ELF2X motif insertion at target promoters in cells modified by PERSIST-On and sorted for GFP expression from a co-transfected plasmid**

Efficiencies of ELF2X insertion in cells that were modified by PERSIST-On (from Fig. 1i) and sorted for GFP expression from a co-transfected plasmid were assessed at various time points post-transfection. Insertion frequencies were assessed for up to 31 days post-transfection at the *IL2RA* and *MYOD1* promoters in K562 cells and at *EPCAM* and *MYOD1* promoters in HEK293T cells. Data shown are three biological replicates and the line represents the mean of those data.

**Extended Data Figure 6: Fine-tuning of PERSIST-On activation of the *IL2RA* and *HER2* gene promoters by systematically mutating EndoTF motifs**

**a-b**, Enrichment scores of a large series of systematically mutated ELF2X (**a**) or CREB/ATF/3-2X (**b**) motifs (blue bars) inserted into the *HER2* promoter in K562 cells that had been flow sorted based on HER2 surface expression as illustrated in Fig. 3a. Mutated motifs are denoted by “base position - base changed nucleotide”. Enrichment scores are also shown for the original unmutated ELF2X or CREB/ATF/3-2X motifs (red bars) and five random negative control DNA sequences (grey bars). Closed circles indicate biological replicates (n=3) and bars represent the means of these replicates.

**c**, Individual validation of ten mutated ELF2X motifs (enriched in the MOSAIC experiments of Fig. 3b) inserted at the *IL2RA* promoter in K562 cells using CRISPR prime editing. An unmutated ELF2X (as a positive control - Pos) and a random DNA sequence (as a negative control - Neg) were also assessed. Top panel shows fold-activation of *IL2RA* gene expression observed and bottom panel shows the efficiency of motif introduction in the same populations of transfected cells. Circles indicate three biological replicates and bars represent the mean of those data.

**d**, Individual validation of nine mutated CREB/ATF/3-2X motifs (enriched in the MOSAIC experiments of Fig. 3c) inserted at the *IL2RA* promoter in K562 cells using CRISPR prime editing. An unmutated ELF2X (as a positive control - Pos) and a random DNA sequence (as a negative control - Neg) were also assessed. Top panel shows fold-activation of *IL2RA* gene expression observed and bottom panel shows the efficiency of motif introduction in the same populations of transfected cells. Circles indicate three biological replicates and bars represent the mean of those data.

**e-f**, Enrichment scores of a large series of systematically mutated ELF2X (**e**) or CREB/ATF/3-2X (**f**) motifs (purple bars) inserted into the *HER2* promoter in K562 cells that had been analyzed by ChIP as illustrated in Fig. 3a. Mutated motifs are denoted by “base position - base changed nucleotide”. Enrichment scores are also shown for the original unmutated ELF2X or CREB/ATF/3-2X motifs (red bars) and five random negative control DNA sequences (grey bars). Closed circles indicate biological replicates (n=3) and bars represent the means of these replicates.

**g-h**, Scatterplot showing mean ChIP enrichment scores of various mutated ELF2X motifs with an anti-ELF4 antibody (**g**) or CREB/ATF/3-2X motifs with an anti-ATF4 antibody (**h**) at the *IL2RA* (x-axis) and *HER2* (y-axis) promoters (data from **e**, **f**, and **Figs. 3f – 3g**). Correlation coefficients (R) were determined with Spearman’s Rho. Blue dots: mutated ELF2X or CREB/ATF/3-2X motifs; grey dots: negative controls, red dots: unmutated ELF2X or CREB/ATF/3-2X motif.

**i-j**, Scatterplots showing ChIP-based enrichment scores (x-axes) and HER2 surface expression-based (flow sorting) enrichment scores (y-axes) for motifs associated with ELF4 (**i**) or ATF4 (**j**) EndoTFs. Correlation coefficients (R) were determined with Spearman’s Rho. Blue dots: mutated ELF2X or CREB/ATF/3-2X motifs; grey dots: negative controls, red dots: unmutated ELF2X or CREB/ATF/3-2X motif.

**Extended Data Figure 7: PWMs and sequence logos derived from MOSAIC-based large-scale screening of systematically mutated EndoTF motifs**

**a**, ELF2X Sequence logo derived from JASPAR database^14^.

**b**, CREB/ATF/3-2X sequence logo derived from JASPAR database^14^.

**c**, *In cellula*-derived ELF2X position weight matrix (PWM) presented as sequence logos at the *IL2RA* promoter based on IL2RA cell surface expression

**d**, *In cellula*-derived CREB/ATF/3-2X position weight matrix (PWM) presented as sequence logos at the *IL2RA* promoter based on IL2RA cell surface expression

**e**, *In cellula*-derived ELF2X position weight matrix (PWM) presented as sequence logos at the *IL2RA* promoter based on ELF4 binding.

**f**, *In cellula*-derived CREB/ATF/3-2X position weight matrix (PWM) presented as sequence logos at the *IL2RA* promoter based ATF4 binding.

**g**, *In cellula*-derived ELF2X position weight matrix (PWM) presented as sequence logos at the *HER2* promoter based on HER2 cell surface expression.

**h**, *In cellula*-derived CREB/ATF/3-2X position weight matrix (PWM) presented as sequence logos at the *HER2* promoter based on HER2 cell surface expression.

**i**, *In cellula*-derived ELF2X position weight matrix (PWM) presented as sequence logos at the *HER2* promoter based on ELF4 binding.

**j**, *In cellula*-derived CREB/ATF/3-2X position weight matrix (PWM) presented as sequence logos at the *HER2* promoter based ATF4 binding.

## ONLINE METHODS

### Plasmids and oligonucleotides

Sequences for pegRNAs and sgRNAs can be found in **Supplementary Table 2**. PegRNA expression plasmids were constructed by ligating previously annealed oligo duplexes consisting of a spacer (target), 3’ extension, and sgRNA scaffold, into BsaI-digested pU6-pegRNA-GG-Vector plasmid (Addgene #132777) as previously described^6^. sgRNA plasmids were constructed by ligating annealed oligo duplexes carrying spacer sequences into BsmBI-digested BPK1520 (Addgene #65777).

### Analysis of EndoTF motifs enriched in active promoters

Promoter regions (+/-250 bp from the TSS) of top-ranked 1000 highly expressed genes and bottom-ranked 4000 lowly expressed genes by FPKM values of HEK293T RNA-seq data (GSM 3700993, GSM3724622)^15,16^ and K562 RNA-seq data (GSM4133283, GSM4133284)^9^ were used for the TF motif enrichment analysis using the Haystack software^3^.

### EndoTF motif selection for library screening

EndoTF motif clusters (n=286) were derived from https://resources.altius.org/~jvierstra/projects/motif-clustering/releases/v1.0/cluster_viz.html. For each TF cluster, a consensus motif was constructed based on the most likely nucleic acid at each position based on the PWM model. For a uniform insertion of 24 bp sequence at the corresponding promoter of target genes, each TF cluster consensus motif was repeated until it was 24 base pairs in length. (**Supplementary Table 1**). In addition to the 286 TF clusters, a positive control (ELF2X) and five randomly generated 24 base pair negative control sequences were added to the TF motif library.

### PCR-generated prime editing guide RNAs

PegRNAs and ngRNAs used for tiling of ELF2X motif insertion (**Extended Data Fig. 1**), EndoTF pooled libraries (Fig. 2) and mutated TF libraries (Fig. 3) were generated by two sequential PCR steps. A detailed protocol is available in **Supplementary Note 1**. Briefly, in the first PCR step, spacer sequence oligos, sgRNA scaffold sequence oligos, and 3’ extension sequence oligos for pegRNAs and spacer sequence oligos and sgRNA scaffold sequence oligos for ngRNAs were assembled. For pegRNA libraries, 3’ extension sequence oligos encoding different TF or mutated TF motifs were pooled together. In the second PCR step, the product from the first PCR was fused to an U6 promoter sequence to generate pegRNA or ngRNA constructs capable of expression in human cells. Reaction and cycling conditions for the PCR steps are given in Supplementary Note 1. The PCR products were cleaned up using paramagnetic beads (0.7-1.2X beads to sample ratio) and 75% ethanol washes.

### Human cell culture conditions

ATCC STR-authenticated HEK293T (CRL-3216) and K562 (ATCC CCL-243) cell lines were used in this study (HEK293T cells were authenticated Dec 13, 2018, and K562 cells were authenticated Nov 8, 2019). All cell culture reagents were obtained from Thermo Fisher unless otherwise specified. HEK293T cells were grown in Dulbecco’s Modified Eagle Medium and K562 cells in Roswell Park Memorial Institute 1640 medium with additional 2 mM Glutamax, supplemented with 10% heat-inactivated fetal bovine serum and 100 units/ml penicillin and 100 µg/ml streptomycin, at 37° C, in 5% CO_2_. Media supernatants were analyzed biweekly for any contamination of the cultures with mycoplasma using MycoAlert PLUS Mycoplasma Detection Kit (Lonza).

### Cell transfections

To screen pegRNAs and ngRNAs for ELF2X insertion efficiencies at different promoters (**Extended Data Fig. 1)**, HEK293T cells were seeded at 1.25 × 10^4^ cells per well in 96-well flat bottom cell culture plates (Corning). After 24 h, cells were transfected with 40 ng of prime editor plasmid and 13.3 ng of pegRNA and 4.4 ng ngRNA using 0.6 µl of TransIT-X2. To characterize the ELF2X motif insertion by the 5 best performing pegRNAs (**Extended Data Fig. 2**) and to test durability of target gene expression (Fig. 1c) in HEK293T cells, cells (6.25-6.8 x 10^4^/well) were seeded in 24-well cell culture plates (Corning) for 24 hours (h) prior to transfection. Cells were lipofected with 300 ng of prime editor plasmid (Addgene #132775), 100 ng of pegRNA plasmid, and 33.3 ng of ngRNA plasmid using 3 µl of TransIT-X2 (Mirus). Cells for CRISPRa experiments, were lipofected with 375 ng of dCas9-VPR plasmid (Addgene #179299) and 125 ng of sgRNA plasmid using 3µl of TransIT-X2. For the enrichment method using GFP plasmid (Lonza) in Fig. 1i, we used the same amount of DNA for PE3 or CRISPRa system as described above and added 16.6 ng of GFP plasmid. To characterize the ELF2X motif insertion by the 5 best performing pegRNAs in K562 cells, (**Extended Data Fig. 2),** cells were seeded at 1.6 x 10^5^ cells/mL medium in a 15 cm-dish and after 24 hours, cells (2 x 10^5^) were nucleofected with the plasmids using a 4D-Nucleofector (Lonza) and the FF-120 program with the SF Cell Line Nucleofector Kit. For experiments to test the durability of PERSIST-On shown in Fig.1d-1e, 1 x 10^6^ cells were nucleofected using 3840 ng prime editor plasmid, 960 ng pegRNA plasmid, and 398.4 ng ngRNA plasmid using the Cell Line Nucleofector Kit V (Lonza). To test the durability of CRISPRa as shown in **Figs. 1d-1e**, 1 x 10^6^ cells were nucleofected using 3750 ng of dCas9-VPR and 1250ng of sgRNA plasmid using the Cell Line Nucleofector Kit V (Lonza). For the enrichment method using GFP plasmid (Lonza), we used the same amount of DNA for PE3 or CRISPRa system as described above and added 199.2 ng of GFP plasmid. To screen different TF libraries or mutated TF libraries for activation of gene expression using prime editing (Figs. 2 and 3), K562 cells (1 × 10^6^) were nucleofected twice with 6000 ng prime editor plasmid, 1500 ng pegRNA pooled libraries, and 500 ng ngRNA, using the SF Cell Line Nucleofector X Kit L (Lonza). Second nucleofection was performed 72 hours after the initial reaction.

### Quantitative reverse transcription PCR

Total RNA was extracted from the cells using the NucleoSpin RNA Plus Kit (Clontech) and 250 ng of purified RNA was used for cDNA synthesis using High-Capacity RNA-to-cDNA Kit (Thermo Fisher Scientific). Expression of the target genes was determined using the cDNA, by quantitative PCR (qPCR) using Fast SYBR Green Master Mix (Thermo Fisher Scientific) and 1 µM gene-specific primers (**Supplementary Table 3)**, in 384-well plates on a LightCycler 480 (Roche) with the following program: initial denaturation at 95 °C for 20 seconds (s) followed by 45 cycles of 95 °C for 3 s and 60 °C for 30 s. Since Ct values fluctuate for transcripts expressed at very low levels, values greater than 35 were considered as 35, and used as the baseline Ct value. Gene expression levels were normalized to *HPRT1* and calculated relative to that of the negative controls (prime editor with non-targeting pegRNAs/ngRNA or dCas9-VPR with non-targeting gRNA). *HPRT1* qPCR control was independently assayed for each sample. Frequency, mean, and standard error of the mean were calculated using GraphPad Prism 8.

### Flow Cytometry

For experiments involving the expression of cell-surface proteins (Fig. 1), cells were washed with cell staining buffer (Biolegend) 72 hours post-transfection and incubated with Phycoerythrin (PE) conjugated IL2RA, HER2 or EpCAM antibodies (Biolegend) for 15 minutes, followed by two washes with cell staining buffer according to the manufacturer’s protocol. All of the PE positive cells for IL2RA, HER2 or EpCAM were sorted using FACS Aria Fusion (BD Biosciences) (Supplementary Note 1) and then passaged every 3 days for testing the durability of expression of the target protein. Protein expression was measured by LSR FortessaX-20 flow cytometer (BD Biosciences). When the cells were transfected with GFP plasmid (Fig. 1i), top 50% of GFP positive cells were sorted and maintained for measuring durability target gene expression. To screen different TF or mutated TF motifs for target gene activation (Figs. 2 and 3), dual-nucleofected K562 cells were stained with IL2RA or HER2 antibodies 72 hours post-second nucleofection and subjected to fluorescence assisted cell sorting to collect cells that were positive and negative for antibody staining, followed by lysing for gDNA extraction.

### Chromatin Immunoprecipitation (ChIP)

ChIP assays for identifying ELF4 recruitment at ELF2X insertion sites with active promoter chromatin state (Fig. 1) were carried out as previously described^9^, using cells (5 x 10^6^) transfected with prime editing components and sorted for IL2RA, HER2, and EpCAM and control cells (5 x 10^6^) that were transfected with prime editor plasmid and non-targeting pegRNA and ngRNA. For ELF4 and H3K4me3 ChIP, 6.6 µg/mL of ELF4 antibody (F-11, Santa Cruz Biotechnology) and 2.3 µg/mL of H3K4me3 antibody (C42D8, Cell Signaling Technology) respectively were used. At sites where the mutated ELF2X or CREB/ATF/3-2X motif libraries were installed (Fig. 3), ELF4 or ATF4 ChIP was performed using double-nucleofected K562 cells as described above with some modifications; 1 x 10^8^ cells were used for this assay, sonicated chromatin was incubated with 33 µg/mL ELF4 antibody (F-11, Santa Cruz Biotechnology) or 11 µg/mL ATF4 antibody (D4B8, Cell Signaling Technology). Antibody-chromatin complexes were pulled down with 30 µl or 100 µl of protein G-Dynabeads (Thermo Fisher Scientific) for ChIP with 5 x 10^6^ and 1 x 10^8^ cells, respectively. DNA was purified with paramagnetic beads as described previously and quantified using Qubit 4 Fluorometer (Thermo Fisher Scientific).

### ChIP-qPCR

The immunoprecipitated DNA was analyzed by qPCR using Fast SYBR Green Master Mix (ThermoFisher Scientific) with the primers (0.2 µM) listed in **Supplementary Table 3** on a LightCycler 480 (Roche) with the following program: initial denaturation at 95 °C for 20 seconds (s) followed by 45 cycles of 95 °C for 3 s and 60 °C for 30 s. Relative enrichment for each target was calculated by normalization to input control.

### Genomic DNA extraction

Cells were lysed overnight by shaking at 55 °C with 905 µl of gDNA lysis buffer (100 mM Tris-HCl at pH 8, 200 mM NaCl, 5 mM EDTA, 0.05% SDS) supplemented with 70 µl of 20 mg ml^−1^ Proteinase K (NEB) and 25 µL of 1 M DTT (Sigma) per 1 x 10^6^ HEK293T or K562 cells. Subsequently, gDNA was extracted from the lysates using 1–2× paramagnetic beads (0.7-1.2X beads to sample ratio), washed twice with 70-80% ethanol, and eluted in 500 µl of 0.1× EB buffer.

### Targeted amplicon sequencing

Libraries for sequencing were prepared by PCR in two steps (PCR1 and PCR2). In PCR1, target sites were amplified using primers that contained Illumina adaptor sequences. The reactions contained 50-100 ng of gDNA, 500 nM each of forward and reverse primer, 200 µM dNTPs, 1 unit of Phusion DNA Polymerase (NEB) and 1X Phusion HF buffer in a total volume of 50 µl. The cycling conditions for PCR1 were 98 °C for 2 min followed by 35 cycles of 98 °C for 10 s, 65°C for 12s and 72 °C for 12s, and a final 72 °C extension for 10 min. PCR1 products were purified using paramagnetic beads (0.7-1.2X beads to sample ratio) according to the amplicon size as described previously and quantified on Qubit 4 Fluorometer (Thermo Fisher Scientific) using 1X DNA high sensitivity kit (Thermo Fisher Scientific). Bead-purified amplicons with Illumina adapters from PCR1 (50-200 ng) were barcoded with Illumina indexes containing sequences complementary to the adapter overhangs in PCR2, using the cycling conditions of 98 °C for 2 min, 7 cycles of 98 °C 10s, 65 °C 30s and 72 °C 30s followed by 72 °C 10 min. The PCR2 products were purified as above and quantified by Qubit 4 Fluorometer. The amplicon libraries thus generated were sequenced with 300 cycles (2 × 150 bp, paired-end) on the Illumina Miseq using 300-cycle MiSeq Reagent Kit v2 (Illumina) or Micro Kit v2 (Illumina). The FASTQ files were downloaded from BaseSpace (Illumina). All the primers used in the above reactions are listed in **Supplementary Table 3.**

### Data analysis

Amplicon sequencing data were analyzed with CRISPResso2 v.2.0.31. Depending on the position and length of the intended edit(s), parameters for CRISPResso2 analysis were modified (-w, -wc, -qwc) to center the quantification window around the intended editing window. Downstream analysis was conducted using Python 3.7.6 with data sourced from ‘Quantification_window_nucleotide_frequency_table.txt’, ‘CRISPResso_quantification_of_editing_frequency.txt’, and ‘Alleles_frequency_table.txt’. Library editing was quantified with the ‘Alleles_frequency_table.txt’ according to inserted motifs.

### Enrichment Score calculation

The number of reads per TF or mutated TF motif was counted and divided by the total read count per sample per replicate. Then, enrichment scores were calculated as the fold-change of samples in positively sorted cells over negatively sorted cells or ChIP over input samples.

### Generating new PWM

*In-cellulo* derived position weight matrices (PWMs) were created with the python package Logomaker. A position probability matrix (PPM) was first derived using logomaker.alignment_to_matrix(to_type = "probability") per sample (positive versus negative sorted cells, ChIP vs input cells) and the matrix was averaged across three replicates. PWM was then calculated with the following formula:

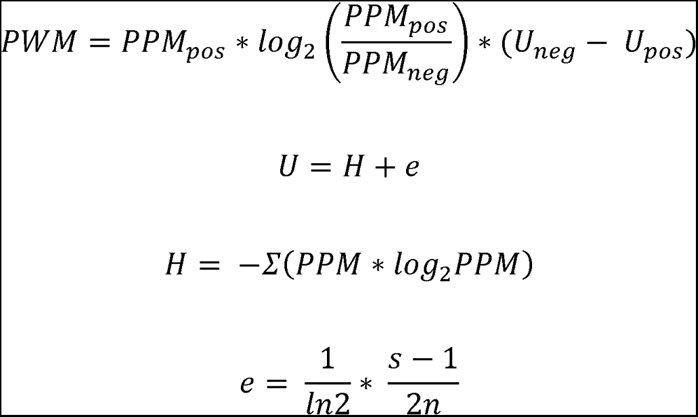

Where *H* denotes entropy and *e* denotes small-sample correction. *s* is 4 for the number of nucleotides and *n* is the number of sequences in the alignment. PPM_pos_ indicates the PPM for positively sorted or ChIPed cells and PPM_neg_ indicates the PPM for negatively sorted or input cells.

### Statistical Analysis

ChIP-qPCR analyses were conducted using Student’s t-test (two-tailed test, paired). The results were considered statistically significant if the p-value was less than 0.05.

## Acknowledgements

J.K.J. was supported by NIH grants R35 GM118158, R01 CA211707, and R01 CA204954, the Desmond and Ann Heathwood MGH Research Scholar Award, and the Robert B. Colvin M.D. Endowed Chair in Pathology. L.P was supported by the NHGRI grant R35 HG010717. We thank Ligi Paul Pottenplackel for comments on the manuscript. Graphical schematics were created with BioRender.com

## Author contributions

Y.E.T., J.Y.H., L.P., and J.K.J. conceived of and designed experiments. Y.E.T., J.Y.H., H.T.S., J.S., I.G.N., and K.C.L. performed experiments and analyzed data. Y.E.T., J.Y.H, J.S., L.P., and J.K.J wrote the manuscript with input from all the authors. L.P. and J.K.J. supervised the project.

## REFERENCES

1. Amabile, A. et al. Inheritable Silencing of Endogenous Genes by Hit-and-Run Targeted Epigenetic Editing. Cell 167, 219–232.e14 (2016).

2. Nuñez, J. K. et al. Genome-wide programmable transcriptional memory by CRISPR-based epigenome editing. Cell 184, 2503–2519.e17 (2021).

3. Pinello, L., Farouni, R. & Yuan, G.-C. Haystack: systematic analysis of the variation of epigenetic states and cell-type specific regulatory elements. Bioinformatics 34, 1930–1933 (2018).

4. Mathelier, A. et al. JASPAR 2016: a major expansion and update of the open-access database of transcription factor binding profiles. Nucleic Acids Res. 44, D110–5 (2016).

5. Curina, A. et al. High constitutive activity of a broad panel of housekeeping and tissue-specific cis-regulatory elements depends on a subset of ETS proteins. Genes Dev. 31, 399–412 (2017).

6. Anzalone, A. V. et al. Search-and-replace genome editing without double-strand breaks or donor DNA. Nature 576, 149–157 (2019).

7. Vierstra, J. et al. Global reference mapping of human transcription factor footprints. Nature 583, 729–736 (2020).

8. Sherwood, R. I. et al. Discovery of directional and nondirectional pioneer transcription factors by modeling DNase profile magnitude and shape. Nat. Biotechnol. 32, 171–178 (2014).

9. Tak, Y. E. et al. Augmenting and directing long-range CRISPR-mediated activation in human cells. Nat. Methods 18, 1075–1081 (2021).

10. Jolma, A. et al. DNA-binding specificities of human transcription factors. Cell 152, 327–339 (2013).

11. Sahu, B. et al. Sequence determinants of human gene regulatory elements. Nat. Genet. 54, 283–294 (2022).

12. Tsai, S. Q. et al. GUIDE-seq enables genome-wide profiling of off-target cleavage by CRISPR-Cas nucleases. Nat. Biotechnol. 33, 187–197 (2015).

13. Collins, R. L. et al. A cross-disorder dosage sensitivity map of the human genome. Cell 185, 3041–3055.e25 (2022).

14. Castro-Mondragon, J. A. et al. JASPAR 2022: the 9th release of the open-access database of transcription factor binding profiles. Nucleic Acids Res. 50, D165–D173 (2022).

15. Grünewald, J., et al. CRISPR DNA base editors with reduced RNA off-target and self-editing activities. Nat. Biotechnol. 37, 1041–1048 (2019).

16. Grünewald, J., et al. Transcriptome-wide off-target RNA editing induced by CRISPR-guided DNA base editors. Nature 569, 433–437 (2019).

