## Extended Data Figures for "CRISPR PERSIST-On enables heritable and fine-tunable human gene activation"

Extended Data Fig. 1

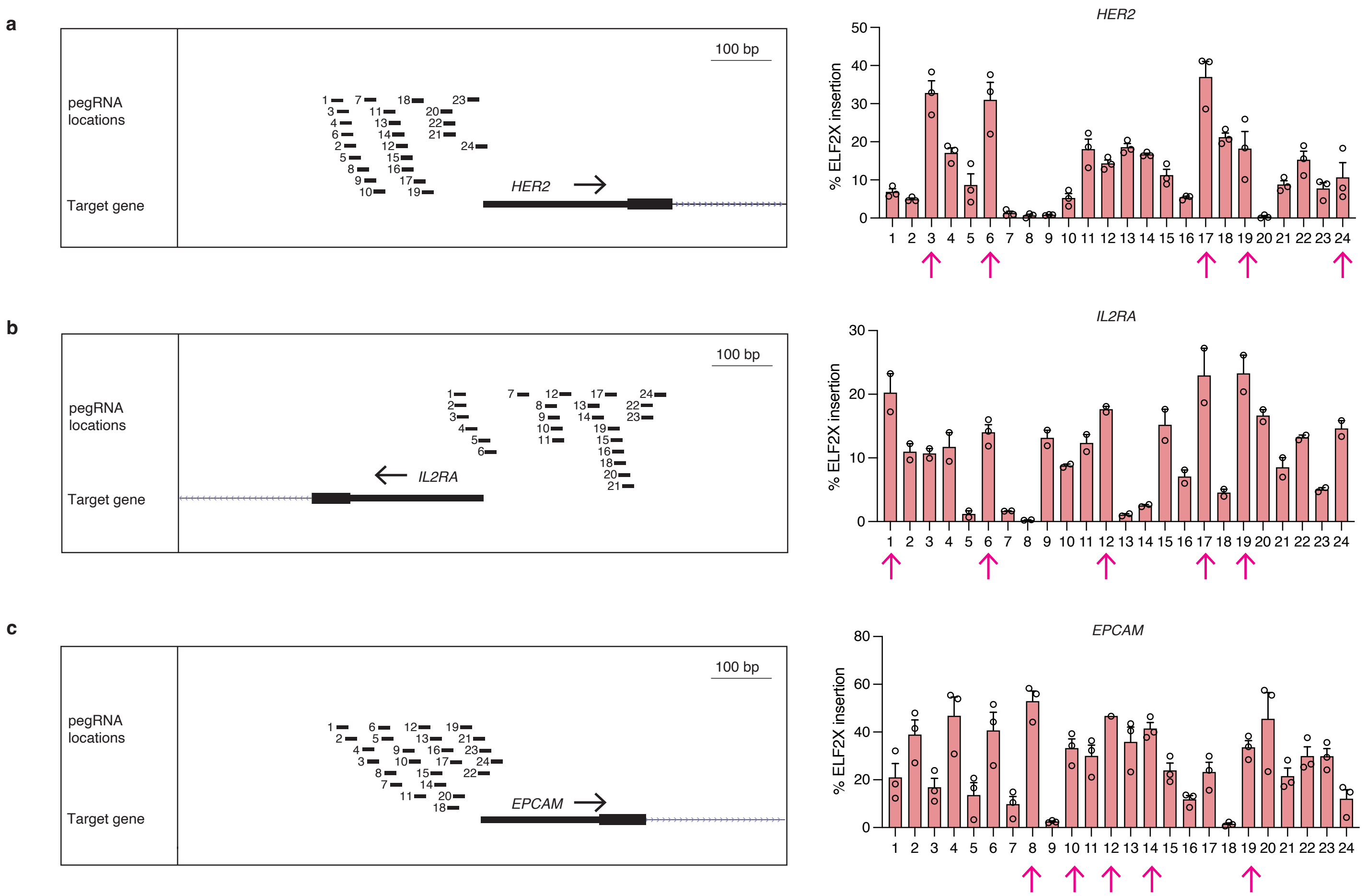

Extended Data Fig. 2

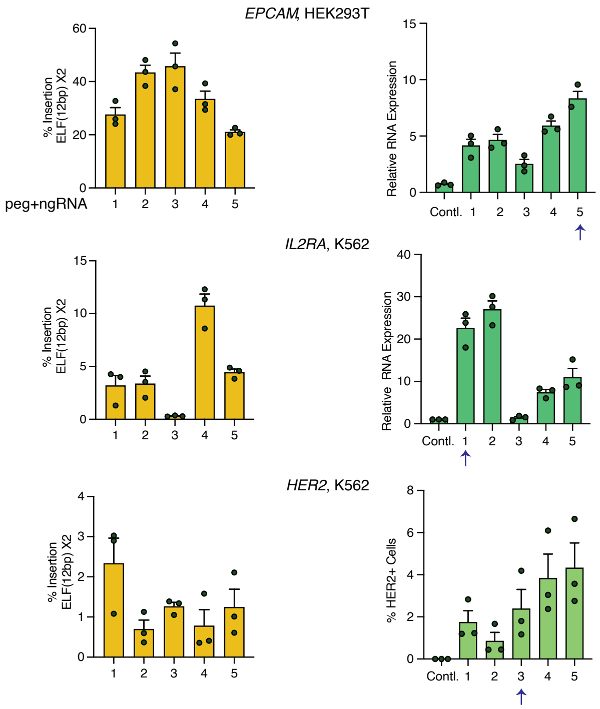

Extended Data Fig. 3

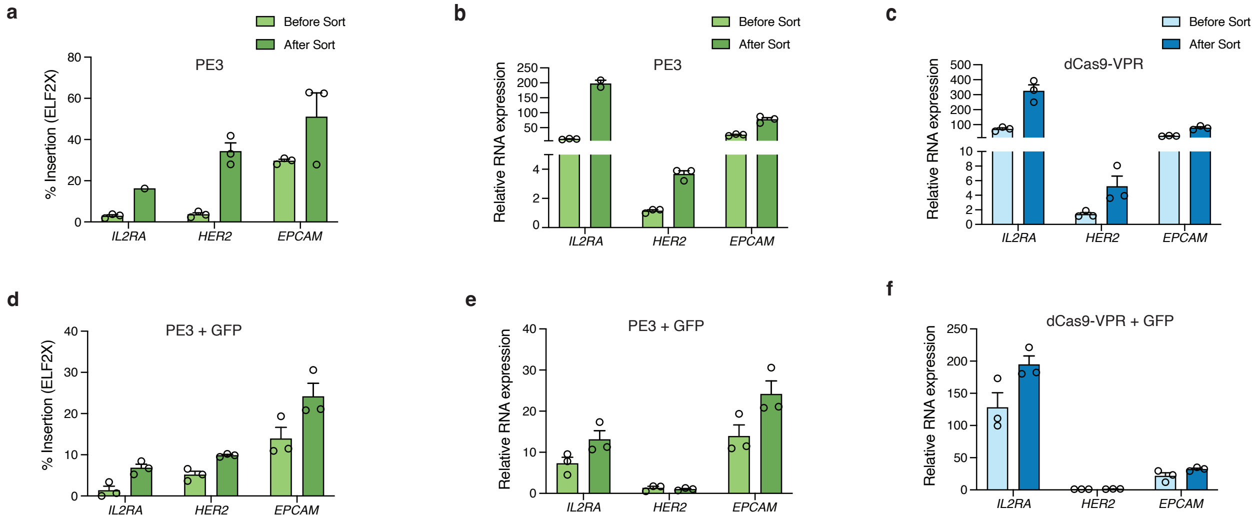

Extended Data Fig. 4

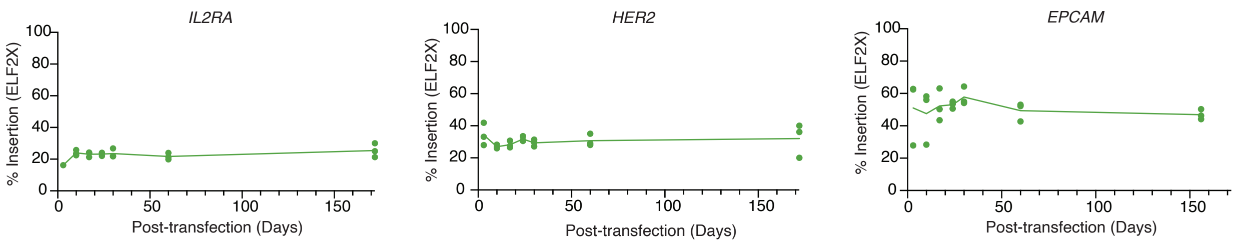

Extended Data Fig. 5

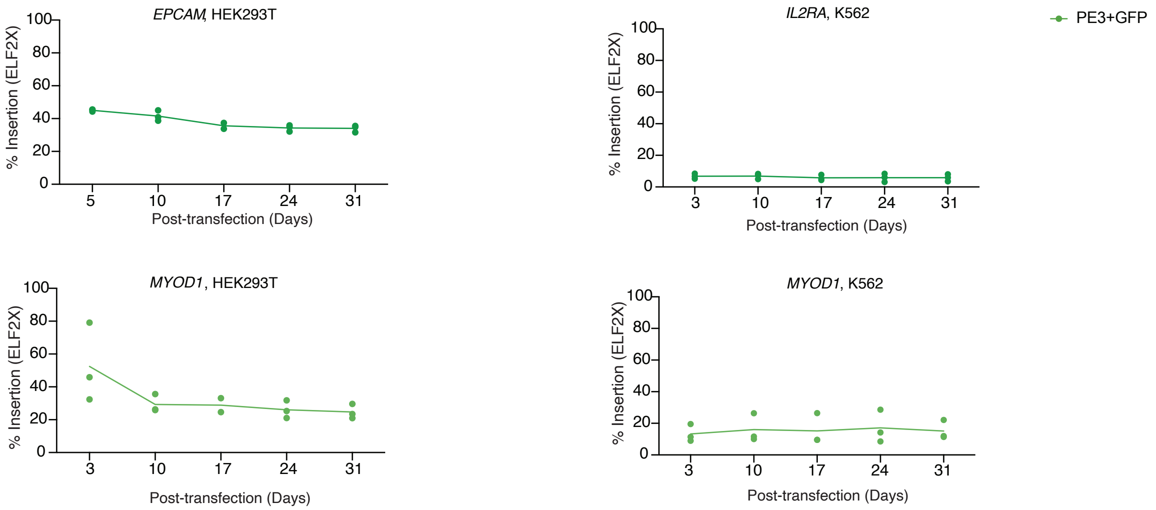

Extended Data Fig. 6

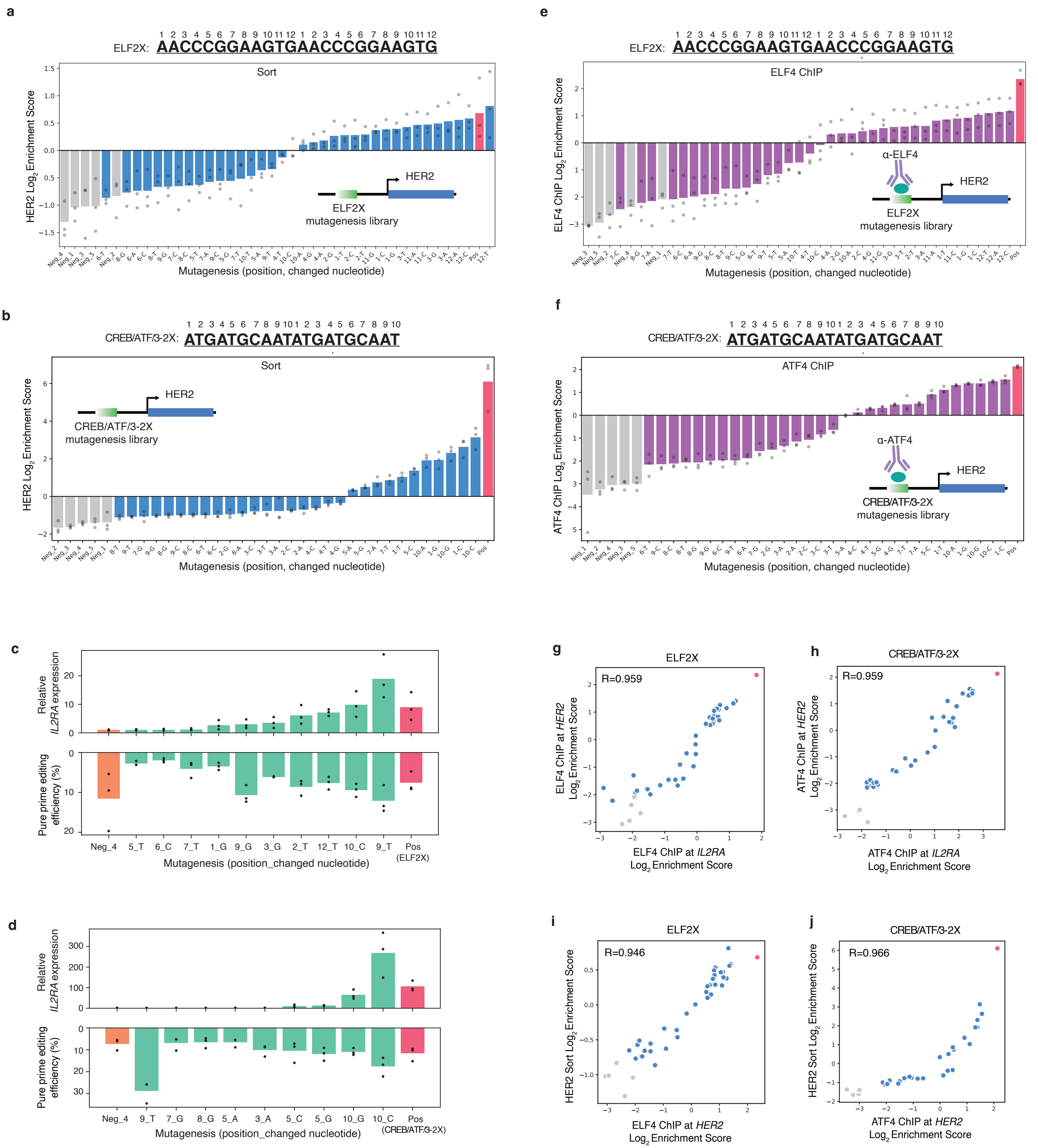

Extended Data Fig. 7

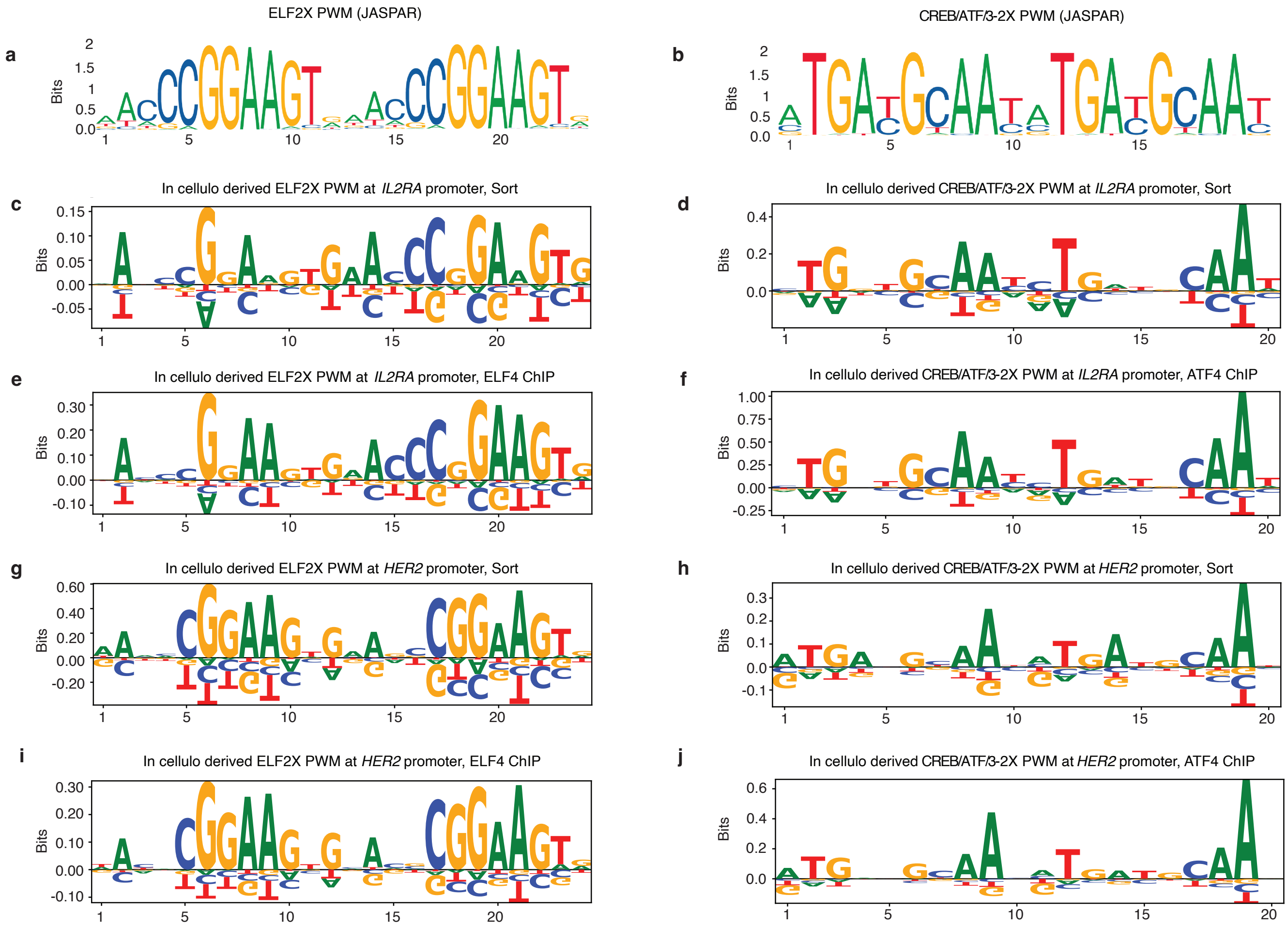
