## Supplementary Note 1 for "CRISPR PERSIST-On enables heritable and fine-tunable human gene activation"

**-Results from Haystack**

**(1)K562**

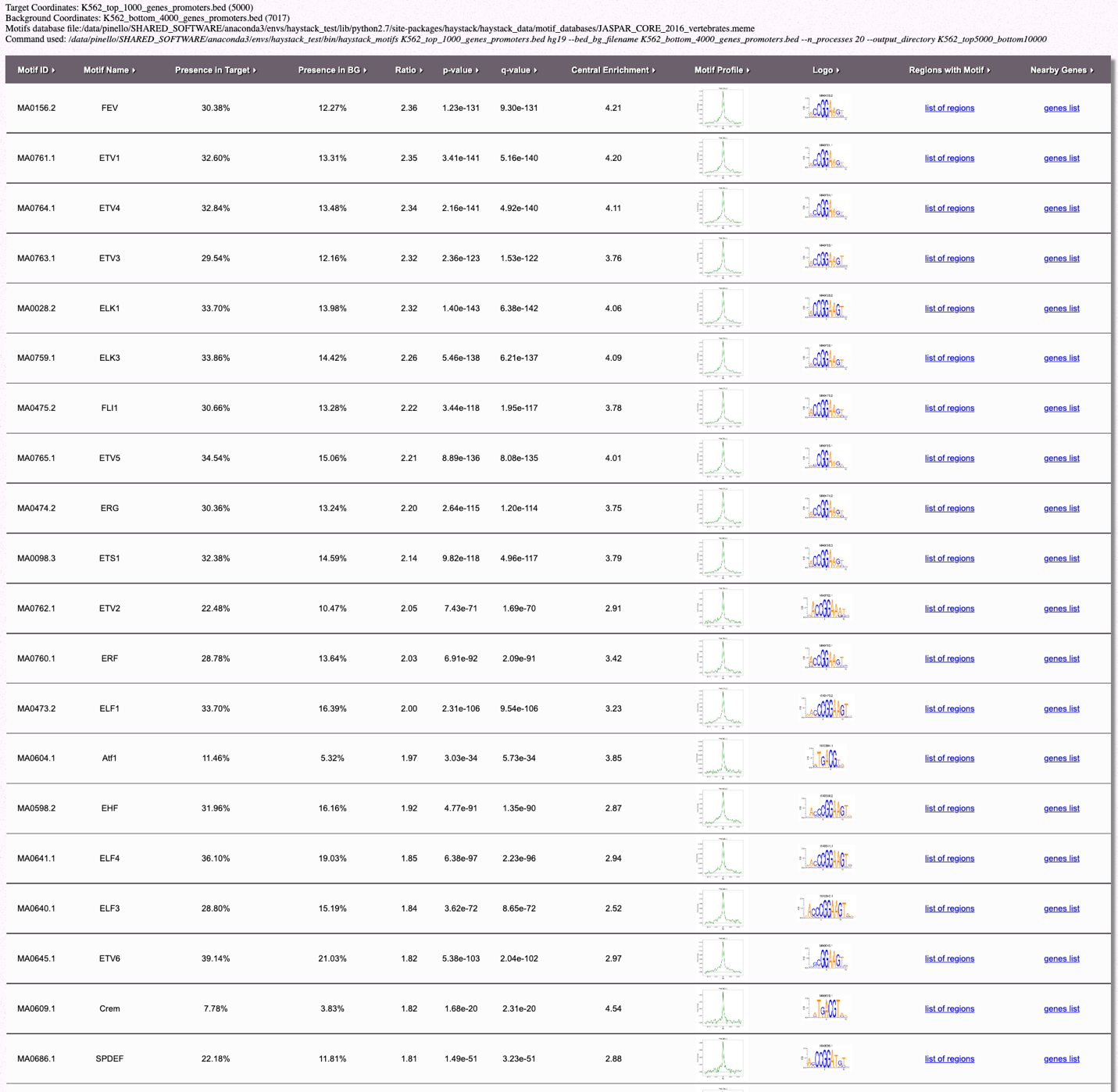

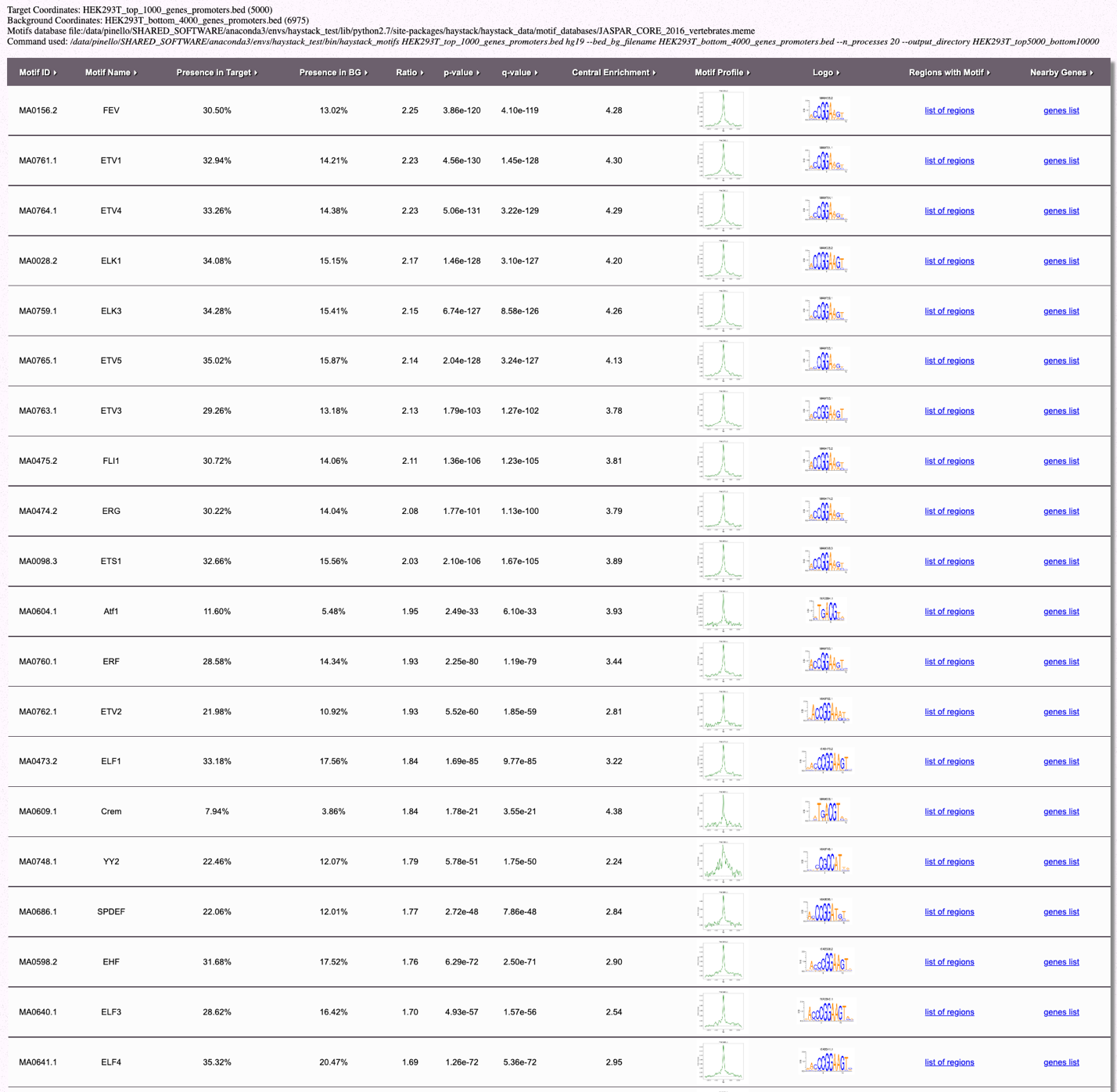
**(2) HEK293T**

**-PCR-generated prime editing guide RNAs (pegRNAs) and nicking guide RNAs (ngRNAs)**

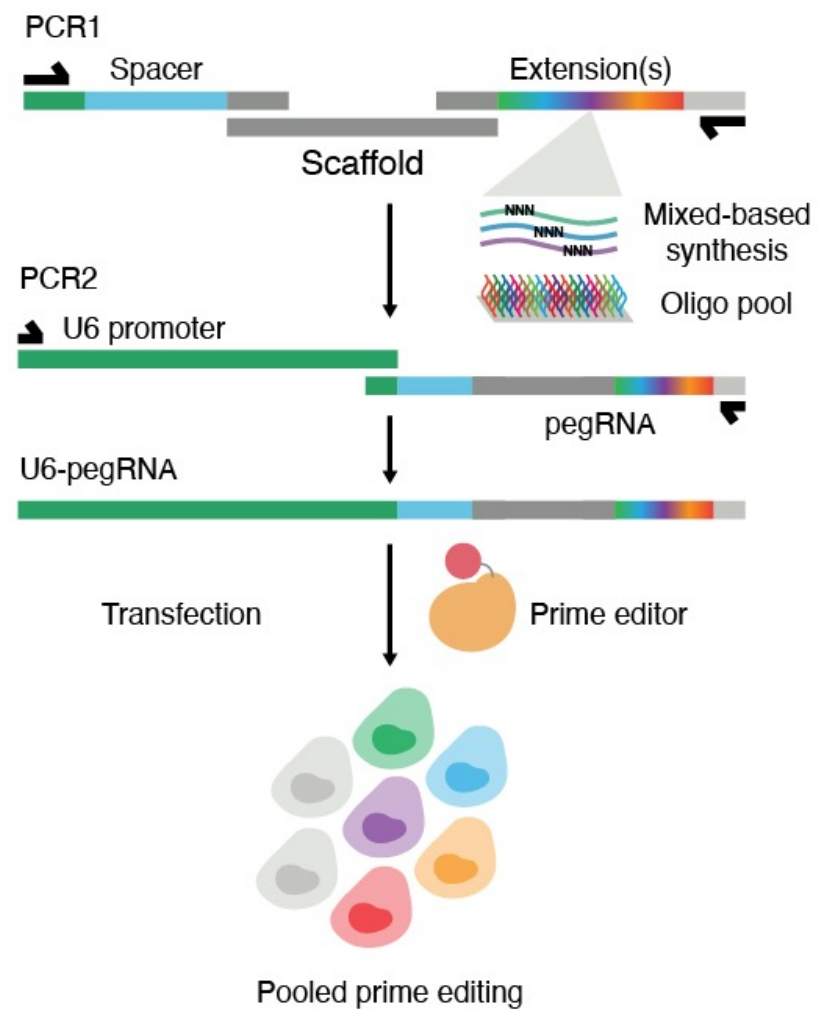

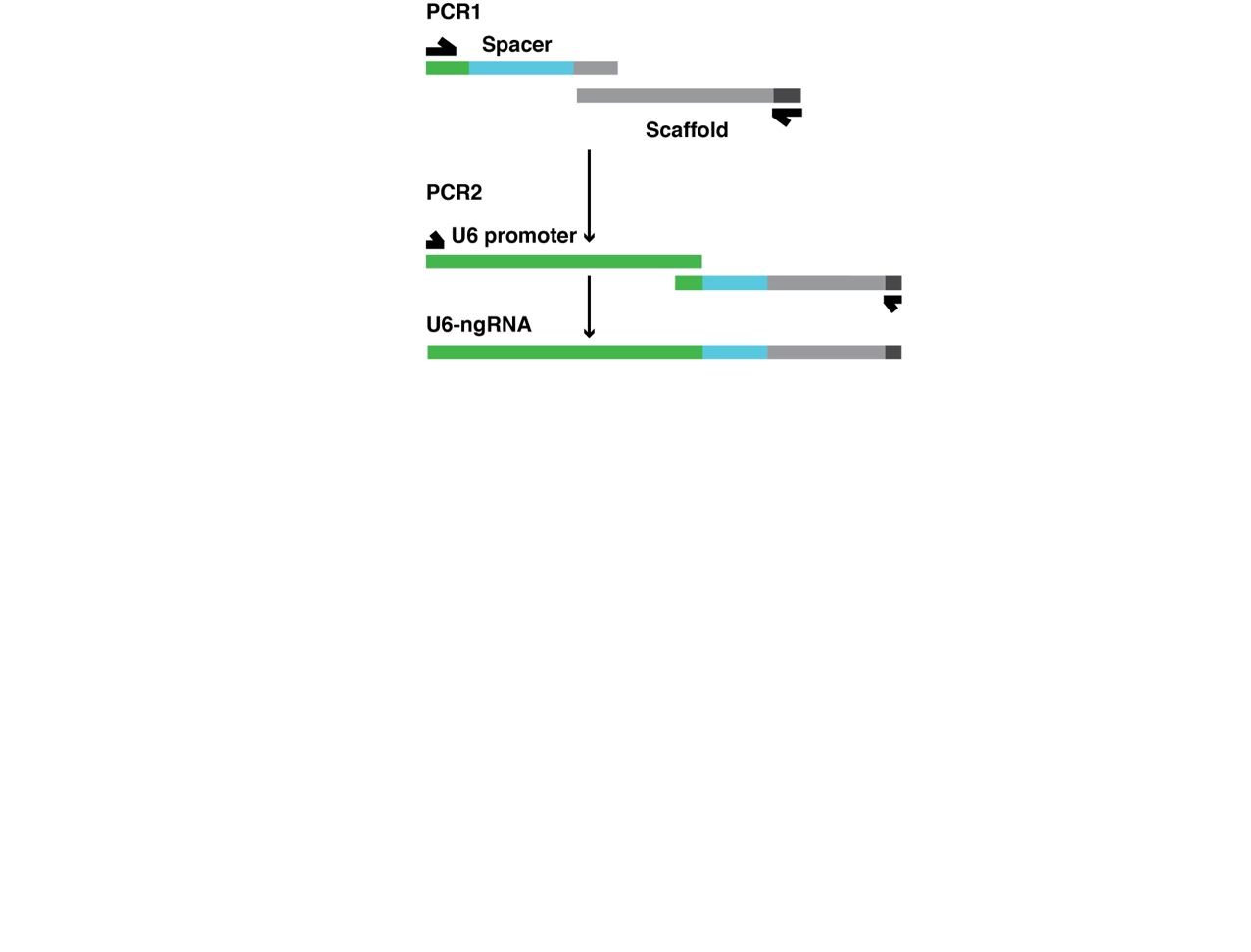

**----------------------------------------------------PCR #1---------------------------------------------------** PCR #1 reaction to Amplify “U6 promoter PCR fragment”

| **Component** | **Volume (µL)** |
| --- | --- |
| 5X PCR Buffer | 10 |
| 10 mM dNTPs | 1 |
| 10 µM U6_Promoter_F | 2.5 |
| 10 µM U6_Promoter_R | 2.5 |
| U6 promoter-containing plasmid (1-5 ng/uL) | 1 |
| Polymerase | 0.5 |
| Water | 32.5 |
| Total | 50 |

lean up with 0.7X paramagnetic beads

PCR1 reaction to assemble “pegRNA PCR fragment”

| **Component** | **Volume (µL)** |
| --- | --- |
| 5X PCR Buffer | 10 |
| 10 mM dNTPs | 1 |
| 10 µM PegRNA_F | 2.5 |
| 10 µM PegRNA_R | 2.5 |
| 10 µM pegRNA_Spacer_Top | 1 |
| 10 µM tracrRNA_Bottom | 1 |
| 10 µM pegRNA_Extension_Top | 1 |
| 10 µM pegRNA_Extension_Bottom | 1 |
| Polymerase | 0.5 |
| Water | 29.5 |
| Total | 50 |

*Clean up with 1.2X paramagnetic beads

PCR1 reaction to assemble “ngRNA PCR fragment”

| **Component** | **Volume (µL)** |
| --- | --- |
| 5X PCR Buffer | 10 |
| 10 mM dNTPs | 1 |
| 10 µM ngRNA_F | 2.5 |
| 10 µM ngRNA_R | 2.5 |
| 10 µM ng_Spacer_Top | 1 |
| 10 µM tracrRNA_Bottom | 1 |
| Polymerase | 0.5 |
| Water | 31.5 |
| Total | 50 |

**-----------------------------------------------------PCR2----------------------------------------------------**

PCR2 reaction to fuse “U6 promoter PCR fragment” with “pegRNA PCR fragment”

| **Component** | **Volume (µL)** |
| --- | --- |
| 5X PCR Buffer | 10 |
| 10 mM dNTPs | 1 |
| 10 µM U6_Promoter_F | 2.5 |
| 10 µM PegRNA_R | 2.5 |
| “U6 promoter PCR fragment” (1-5 ng/uL) | 1 |
| “pegRNA PCR fragment” (1-5 ng/uL) | 1 |
| Polymerase | 0.5 |
| Water | 31.5 |
| Total | 50 |

*Clean up with 0.7X paramagnetic beads

PCR2 reaction to fuse “U6 promoter PCR fragment” with “ngRNA PCR fragment”

| **Component** | **Volume (µL)** |
| --- | --- |
| 5X PCR Buffer | 10 |
| 10 mM dNTPs | 1 |
| 10 µM U6_Promoter_F | 2.5 |
| 10 µM ngRNA_R | 2.5 |
| “U6 promoter PCR fragment” (1-5 ng/uL) | 1 |
| “PegRNA PCR fragment” (1-5 ng/uL) | 1 |
| Polymerase | 0.5 |
| Water | 31.5 |
| Total | 50 |

*Clean up with 0.7X paramagnetic beads

1. Amplicon sequence for TF library and mutagenesis pegRNA PCR product for *IL2RA* promoter (N: Inserted TF motif (24bp for different TF library and ELF2X mutagenesis TF library; 20bp for ATF/CREB 2X mutagenesis TF library))

CTGTACAAAAAAGCAGGCTTTAAAGGAACCAATTCAGTCGACTGGATCCGGTACCAAGGTCGGGCAGGAAGAGGGCCTATTTCCCATGATTCCTTCATATTTGCATATACGATACAAGGCTGTTAGAGAGATAATTAGAATTAATTTGACTGTAAACACAAAGATATTAGTACAAAATACGTGACGTAGAAAGTAATAATTTCTTGGGTAGTTTGCAGTTTTAAAATTATGTTTTAAAATGGACTATCATATGCTTACCGTAACTTGAAAGTATTTCGATTTCTTGGCTTTATATATCTTGTGGAAAGGACGAAACACCGGGATGAGAGAAGAGAGTGCTGTTTTAGAGCTAGAAATAGCAAGTTAAAATAAGGCTAGTCCGTTATCAACTTGAAAAAGTGGCACCGAGTCGGTGCATTGGGCTGGCGTGTTCAGCCAGGAAACTGCCTAGCNNNNNNNNNNNNNNNNNNNNNNNNACTCTCTTCTCTCATTTTTTTGGTGTATCACGTGCTGC

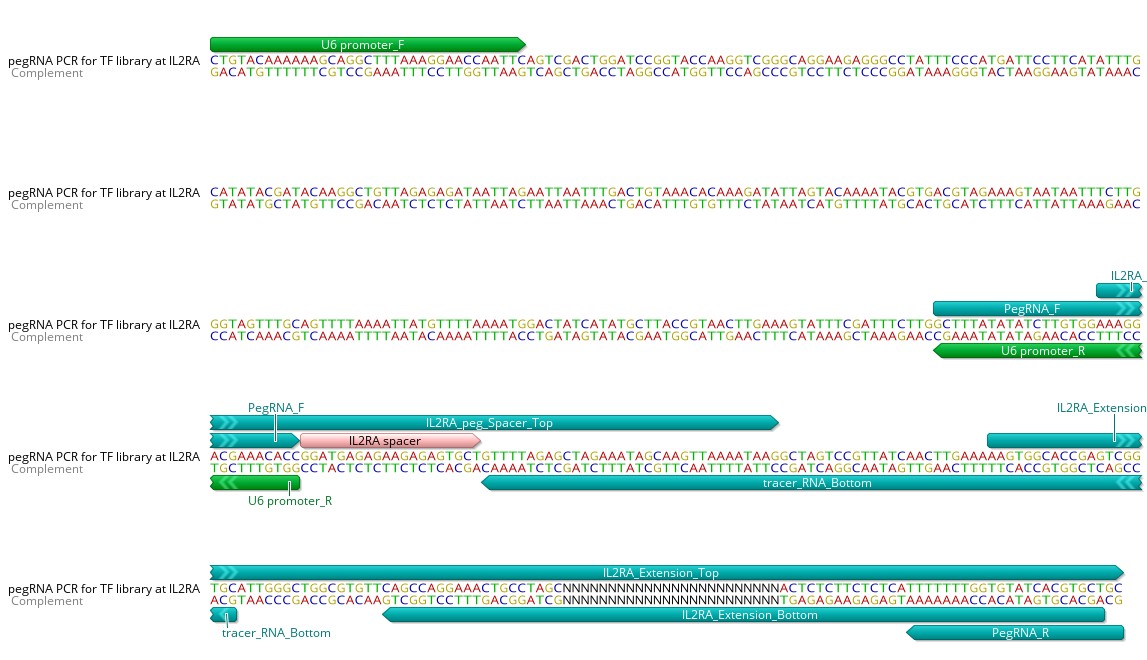

2. Amplicon sequence for TF library and mutagenesis ngRNA PCR product for *IL2RA* promoter

CTGTACAAAAAAGCAGGCTTTAAAGGAACCAATTCAGTCGACTGGATCCGGTACCAAGGTCGGGCAGGAAGAGGGCCTATTTCCCATGATTCCTTCATATTTGCATATACGATACAAGGCTGTTAGAGAGATAATTAGAATTAATTTGACTGTAAACACAAAGATATTAGTACAAAATACGTGACGTAGAAAGTAATAATTTCTTGGGTAGTTTGCAGTTTTAAAATTATGTTTTAAAATGGACTATCATATGCTTACCGTAACTTGAAAGTATTTCGATTTCTTGGCTTTATATATCTTGTGGAAAGGACGAAACACCGTTGATGACAATATAGTTTGGTTTTAGAGCTAGAAATAGCAAGTTAAAATAAGGCTAGTCCGTTATCAACTTGAAAAAGTGGCACCGAGTCGGTGCTTTTTTTGGTGTATCACGTGCTGC

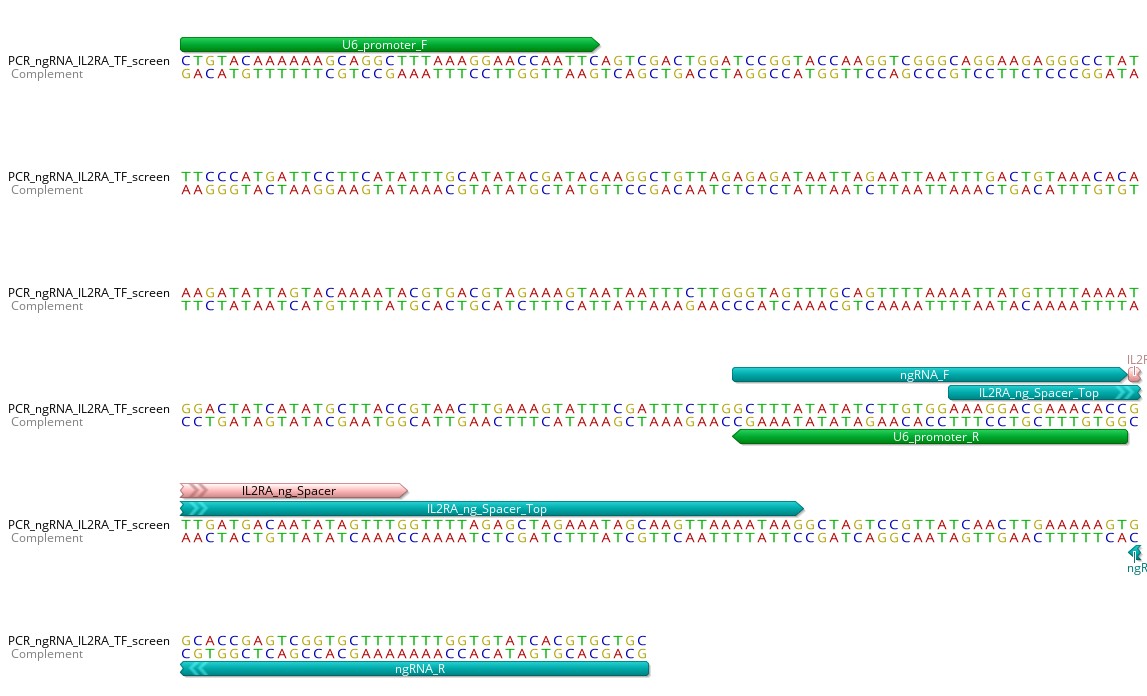

3. Amplicon sequence for TF library and mutagenesis pegRNA PCR product for *HER2* promoter

N: Inserted TF motif (24bp for different TF library and ELF2X mutagenesis TF library; 20bp for ATF/CREB 2X mutagenesis TF library)

CTGTACAAAAAAGCAGGCTTTAAAGGAACCAATTCAGTCGACTGGATCCGGTACCAAGGTCGGGCAGGAAGAGGGCCTATTTCCCATGATTCCTTCATATTTGCATATACGATACAAGGCTGTTAGAGAGATAATTAGAATTAATTTGACTGTAAACACAAAGATATTAGTACAAAATACGTGACGTAGAAAGTAATAATTTCTTGGGTAGTTTGCAGTTTTAAAATTATGTTTTAAAATGGACTATCATATGCTTACCGTAACTTGAAAGTATTTCGATTTCTTG*GCTTTATATATCTTGTGGAAAGGACGAAACACC*GCCCTCTCTTCGCGCAGGCCTGTTTTAGAGCTAGAAATAGCAAGTTAAAATAAGGCTAGTCCGTTATCAACTTGAAAAAGTGGCACCGAGTCGGTGCAGGCGTCCCGGCGCTAGGAGGGACGCACCCAGGNNNNNNNNNNNNNNNNNNNNNNNNCCTGCGCGAAGATTTTTTTGGTGTATCACGTGCTGC
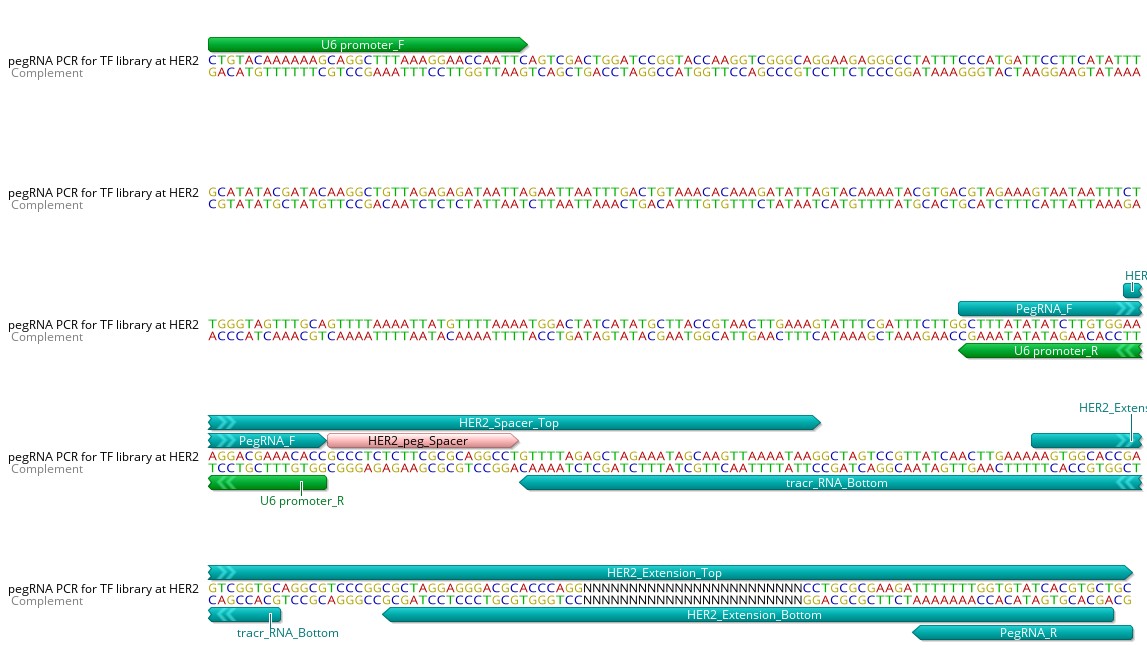

4. Amplicon sequence for TF library and mutagenesis ngRNA PCR product for *HER2* promoter

CTGTACAAAAAAGCAGGCTTTAAAGGAACCAATTCAGTCGACTGGATCCGGTACCAAGGTCGGGCAGGAAGAGGGCCTATTTCCCATGATTCCTTCATATTTGCATATACGATACAAGGCTGTTAGAGAGATAATTAGAATTAATTTGACTGTAAACACAAAGATATTAGTACAAAATACGTGACGTAGAAAGTAATAATTTCTTGGGTAGTTTGCAGTTTTAAAATTATGTTTTAAAATGGACTATCATATGCTTACCGTAACTTGAAAGTATTTCGATTTCTTGGCTTTATATATCTTGTGGAAAGGACGAAACACCGCTGCATTTAGGGATTCTCCGGTTTTAGAGCTAGAAATAGCAAGTTAAAATAAGGCTAGTCCGTTATCAACTTGAAAAAGTGGCACCGAGTCGGTGCTTTTTTTGGTGTATCACGTGCTGC

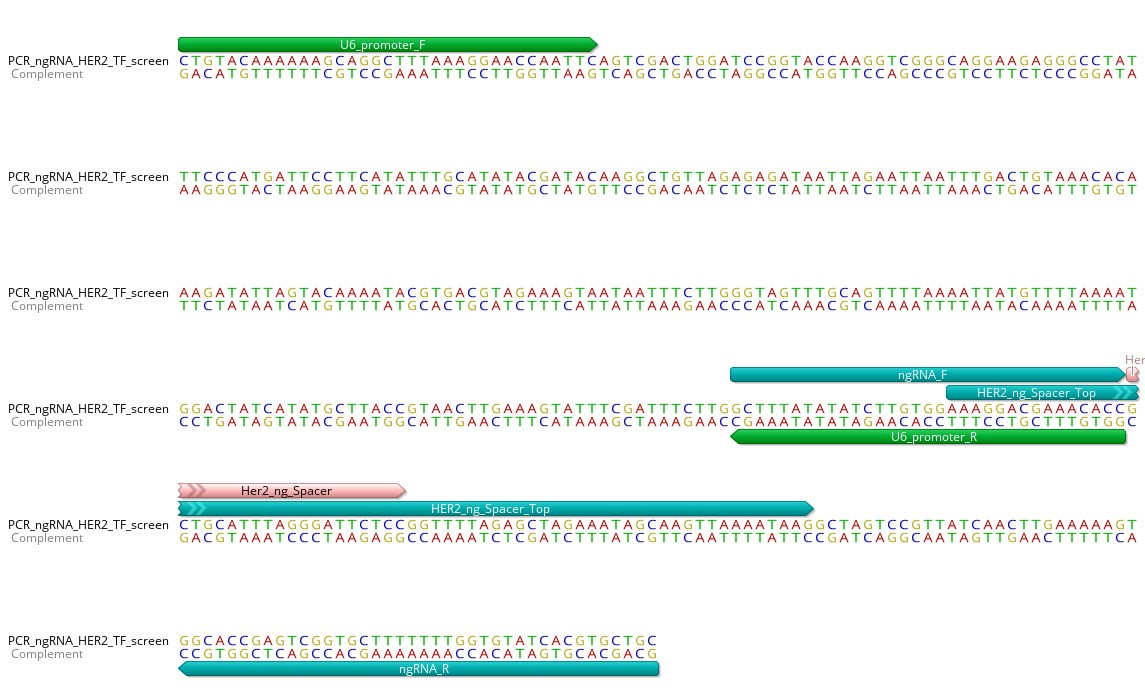

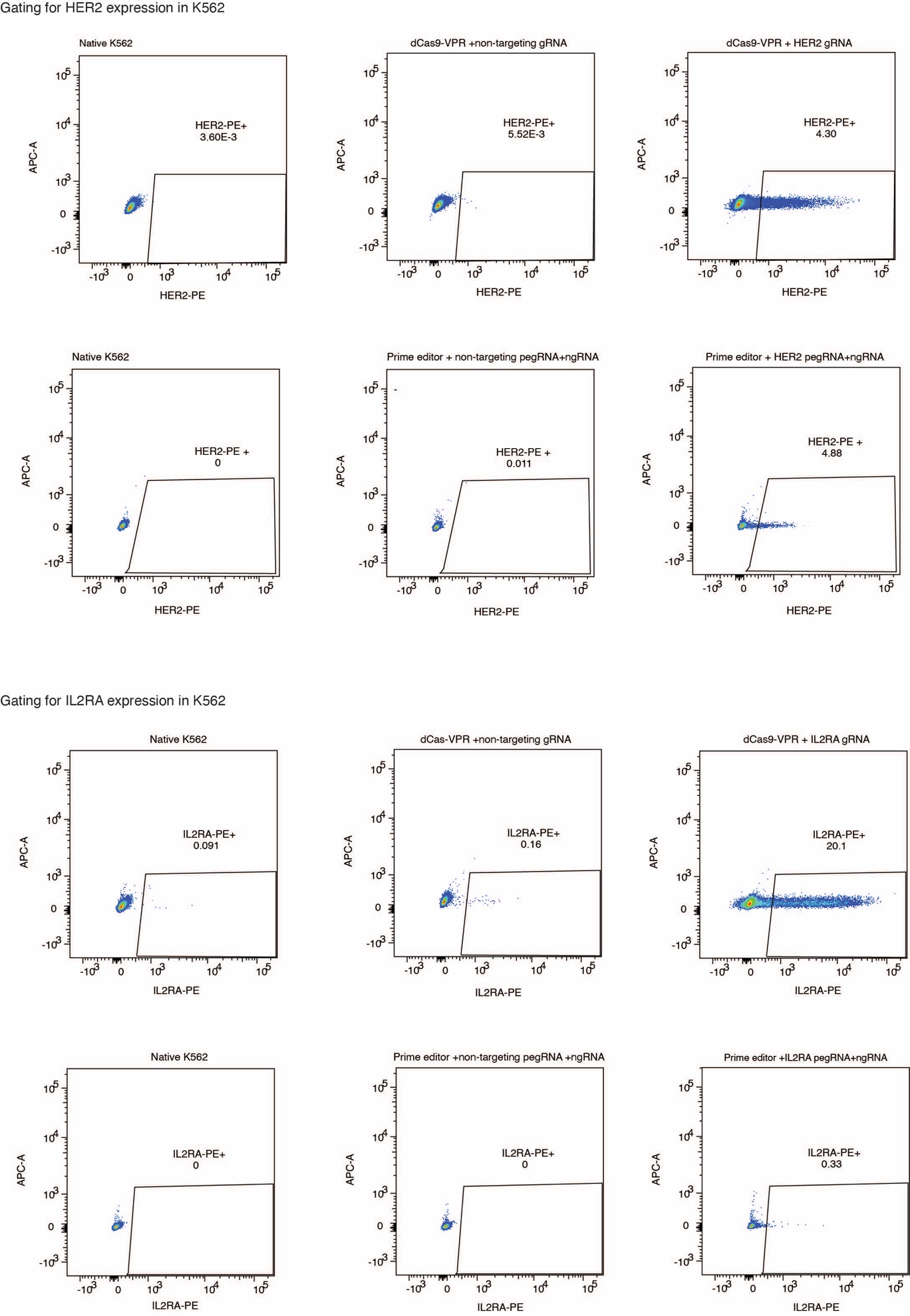

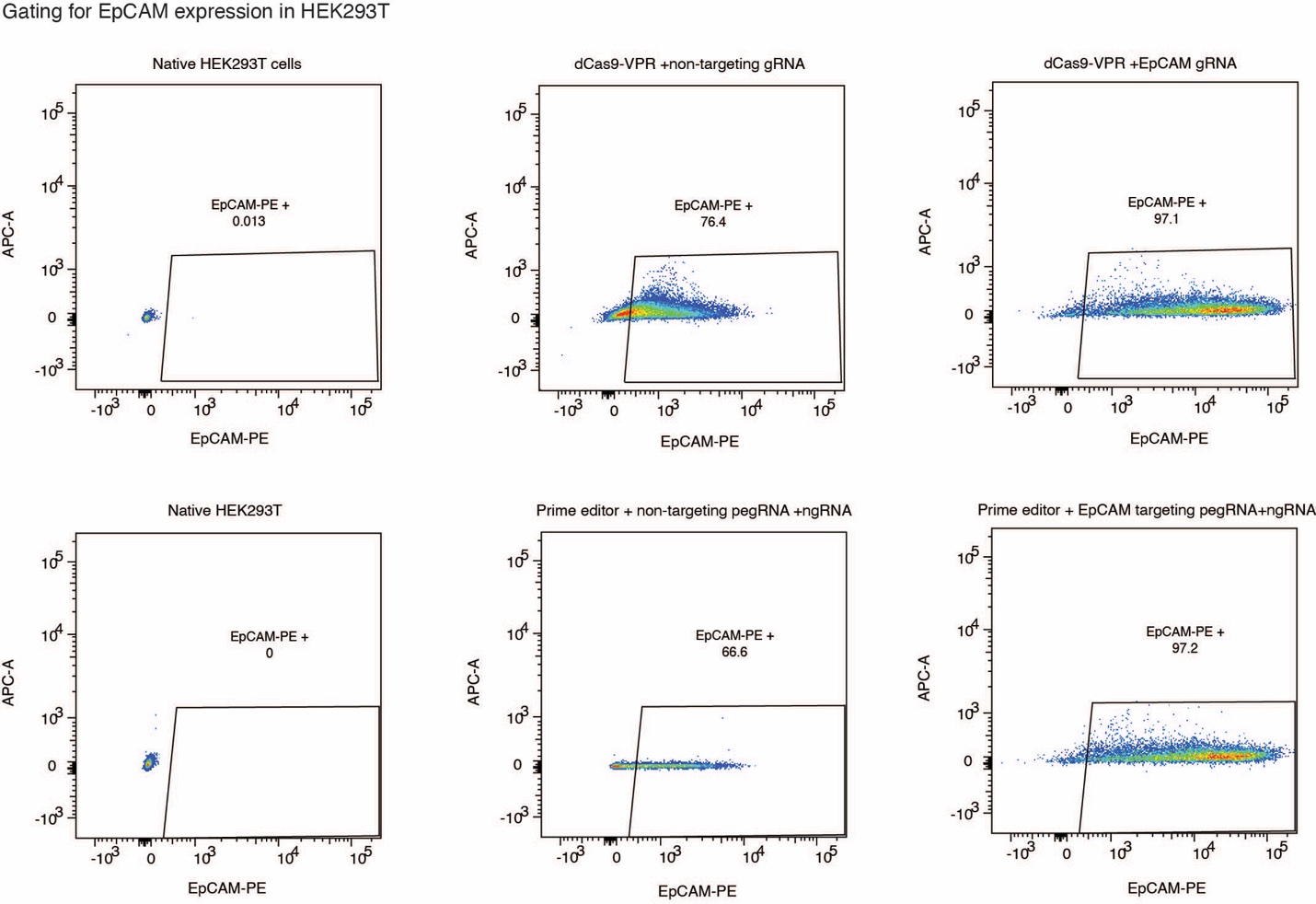
